# Activation of oligonucleotide polyanions using collisions, electrons and photons in a timsOmni platform

**DOI:** 10.1101/2025.11.10.687689

**Authors:** Frédéric Rosu, Rim Chiba, Arjun Mani Mallika, Athanasios Smyrnakis, Jean-François Greisch, Dimitris Papanastasiou, Valérie Gabelica

**Affiliations:** School of Pharmaceutical Sciences, University of Geneva, Geneva, Switzerland; Fasmatech Science & Technology, Athens, Greece; Bruker Switzerland AG, Fällanden, Switzerland

## Abstract

We describe here various ion activation experiments realized in the Omnitrap™ platform integrated on the timsOmni^TM^ mass spectrometer for the analysis of oligonucleotides in the negative ion mode. The activation methods include resonance collision-induced dissociation (_R_CID), electron detachment dissociation (EDD), infrared laser multiple-photon activation (IRMPD) and UV laser photodissociation (UVPD). Special emphasis is given to EDD, either as a standalone technique or in conjunction with vibrational re-activation of the ion radicals. We describe EDD on standard 6-mer DNA sequences that have been extensively characterized on other instruments, followed by a comparison of several activation approaches for the phosphorothioate-based oligonucleotide therapeutics Fomivirsen, and concluding with the fragmentation analysis of 46-mer DNA and RNA. EDD alone already provides excellent sequence information on Fomivirsen, but MS^3^ combinations such as EDD-_R_CID or EDD-IRMPD proved even more effective, including for the 46-mer DNA (less prone to fragmentation than RNA) at a relatively low charge state. The diversity of ion activation combinations available on the Omnitrap platform is demonstrated by an MS^4^ experiment investigating the fate of *a*• and *z*• radical fragments produced by EDD.

**TOC graphics:** 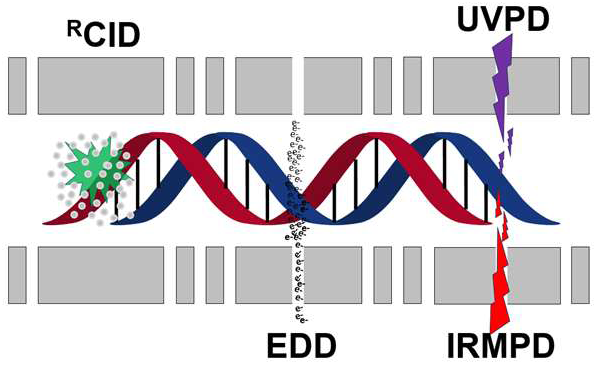

## INTRODUCTION

Top-down sequencing by mass spectrometry entails ionizing intact biopolymers and extensively fragmenting their gas phase ions to localize every residue and associated modification. This approach is gaining momentum for the characterization of protein modification variants, known as proteoforms.^1, 2^ Among ion activation and fragmentation techniques, traditional collision-induced dissociation (CID) typically provides incomplete sequence coverage for intact proteins.^3–5^ A major breakthrough in protein analysis came with the discovery of electron-capture dissociation (ECD).^6^ In ECD, low energy (∼1 eV) electrons are captured by multiply charged protein cations confined within a high-resolution Fourier transform ion cyclotron resonance (FTICR) mass analyzer^7^ or in a linear radiofrequency-free magnetic cell,^8^ forming cation radicals that promote efficient N─Cα bond cleavage along the backbone. A key advantage of ECD is that labile post-translational modifications (PTMs), such as phosphorylations or glycosylations, are preserved while the protein backbone fragments efficiently. Non-covalent interactions, including salt bridges can also be retained during dissociation.^9–12^ An equally powerful technique is electron transfer dissociation (ETD).^13^ In ETD, electrons are transferred from reagent anions to multiply charged protein cations stored simultaneously within three-dimensional or linear quadrupole ion trapping devices, yielding fragmentation behavior analogous to ECD. Besides, other electron-based activation techniques were introduced for peptides and proteins,^14^ for example electron-ionization dissociation (EID), with electron kinetic energies typically > 20 eV.^15^

In the context of oligonucleotide analysis, CID is also limited in terms of sequence coverage. Interpretation becomes increasingly difficult for sequences longer than 20 to 25 nucleotides, as fragment spectra are dominated by base losses and internal fragments rather than the sequence-informative terminal fragments. Nevertheless, achieving comprehensive top-down characterization of intact oligonucleotides is critical for several applications, including the analysis of oligonucleotide therapeutics and their impurities,^16–19^ simultaneous localization of multiple natural modifications^20^, for example in tRNA,^21–24^ mapping of artificial modifications in biophysical studies using chemical labeling,^25–28^ and identification of ligand binding sites in native mass spectrometry experiments.^29–32^

Unlike proteins, nucleic acids ionize more efficiently in negative ion mode, and because electron transfer to multiply charged anions is thermodynamically unfavorable, ECD and ETD, although feasible,^33-^

^35^ are not ideally suited for oligonucleotide fragmentation. When oligonucleotide polyanions are instead irradiated with higher energy electrons, typically ∼16 eV, electron detachment occurs leading to dissociation. This process is known as electron detachment dissociation (EDD).^36–39^ Originally implemented in ultrahigh-vacuum FTICR mass analyzers, EDD has been more recently realized in a plasma-enriched configuration on a custom-modified Sciex ZenoTOF instrument.^40, 41^ Negative ion ETD (NETD), in which radicals are generated by electron transfer from anionic analytes to fluoranthene radical cations, can also induce RNA fragmentation.^42, 43^ Recent NETD studies demonstrated fragmentation of 20-nt RNA followed by activation in the HCD cell (high energy collision) in an orbitrap,^42, 44^ and of miRNA subjected to either CID or IRMPD in an FTICR mass spectrometer.^43^ However, applications to longer oligonucleotide strands have not yet been reported.

Previously, we investigated the irradiation of nucleic acid multiply charged anions with a UV laser in a quadrupole ion trap, using a wavelength range where nucleic acids were known to absorb (∼260 nm). Electron detachment was observed, particularly for strands containing guanines,^45, 46^ revealing a new means of generating radicals from multiply charged anions. Subsequent re-activation of these radical ions by helium resonance CID in an MS^3^ experiment led to efficient backbone cleavage.^45, 47^ This process, coined electron photodetachment dissociation (EPD) or activated electron photodetachment (a-EPD), was later extended to proteins.^48^ The Brodbelt group further explored both a-EPD and direct UV photodissociation of oligonucleotides at 193 nm.^47, 49, 50^ Recent studies of long single guide RNA (∼100-mers) showed that CID alone provides the best sequence coverage (60%),^51, 52^ whereas combining CID with UVPD and a-EPD from independent experiments increased coverage.^53^

The observed product ions depend on the activation method and experimental conditions. The fragmentation nomenclature was established by McLuckey (Figure 1),^54^ and dissociation pathways have been reviewed.^55–57^ Briefly, for negative ions, vibrational activation (via collisions or IR laser irradiation) mainly produce base losses, *a-*Base (*a*-B) and *w* ions for DNA, and *c* and *y* ions for RNA. EDD and UVPD predominantly yield *d* and *w* ions, whereas NETD generates additional *a* and *z* ions, and a-EPD produces *a/w*, *c/y* and *d* ions. Comparisons among activation methods are complicated by the fact that each technique has typically been implemented on different hybrid instrument platforms and applied to distinct analyte classes (e.g., RNA vs. DNA). For instance, in DNA analyzed by a-EPD in helium-filled ion traps, the resulting fragments consisted exclusively of *a*•/*w* and *d/z*• radical ions.^45, 47^ Plasma EDD produced mainly *a*•/*w* and *d/z*•, but also *c* and *x* fragments.^41^ In RNA, a-EPD followed by HCD activation of the radicals species in an Orbitrap Lumos mass spectrometer (Thermo Fisher Scientific) generated predominantly closed shell *w* and *d* ions,^49, 58^ resembling the fragmentation pattern observed in EDD, although plasma EDD additionally yielded small amounts of a• and even fewer z• radical ions.^41^ Tauscher and Breuker previously proposed that *a*• radical ions could rearrange into closed shell *d* ions, and *z*• radical ions to *w* ions.^37^ This in turn suggests that the extent of radical ion survival may reflect the internal energy distribution imparted during EDD. However, the *a*• → *d* and *z*• → *w* reactions have not been experimentally validated.

**Figure 1:**
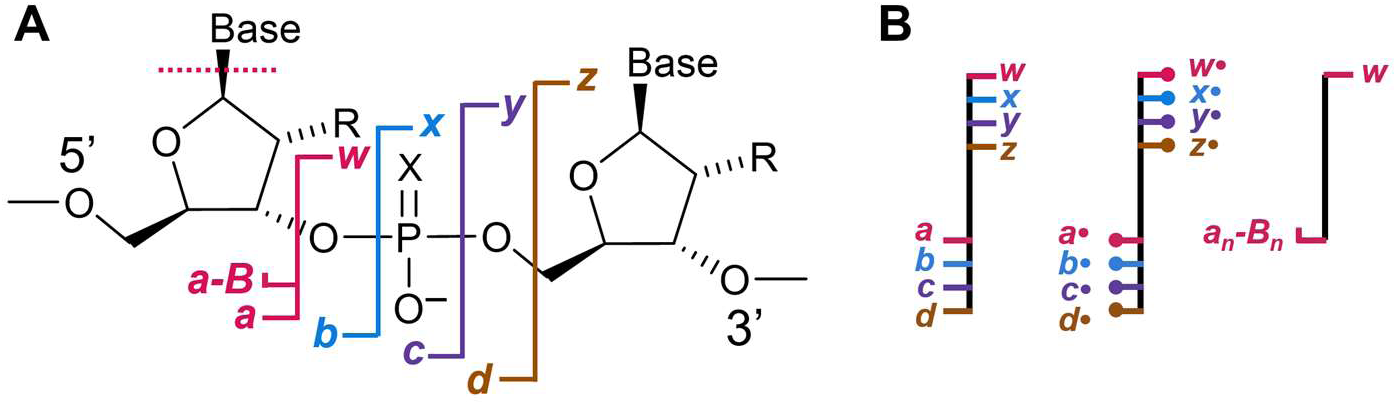
**A**) Nomenclature of oligonucleotide fragments. DNA: R = H, X = O; RNA: R = OH, X = O; phosphorothioate DNA: R = H, X = S. **B**) Schematic annotation of the sequence coverage. The special annotation for *a-*B fragments denotes the specific fragmentation pathway at stake for DNA, where *a/w* fragmentation is triggered by the loss of the base directly adjacent on the 5’-end.

The Omnitrap™ linear ion trap^59^ constitutes an ideal platform to evaluate different activation techniques on ions produced and stored under comparable conditions. Furthermore, several activation methods can be combined in MS^n^ experiments, to improve intact top-down characterization or to explore dissociation pathways. The Omnitrap platform consists of multiple linear ion trap sections arranged in series wherein different fragmentation techniques and MS selection stages combined with intermediate ion enrichment steps can be seamlessly applied. A first section is configured to perform ion isolation (for MS^n^ experiments) and resonance excitation CID, a second section enables ion interactions with electrons whose kinetic energies are tunable in the range 0—100 eV, making it possible to identify optimum conditions for high efficiency EDD experiments, and a third section defines a trapping region where optical access is established for either IR or UV laser access. We report here on the first exploration of a highly diverse range of ion activation methods applied to oligonucleotides on a timsOmni platform, with a special focus on EDD.

## EXPERIMENTAL SECTION

### Sample preparation

6-mer and 46-mer oligonucleotides were purchased from Eurogentec. 46-mers were desalted using Amicon (3 kDa cut-off) ultra centrifugal filter and prepared in 30 mM ammonium acetate (NH_4_OAc 5M stock solution BioUltra grade from Merck). 46-mer oligonucleotide sequences are 5’- TTGCTTAAGTATAAGGATCTAAGTAAAATTTGTCGGTATCTCGGTT-3’ for DNA and 5’- UUGCUUAAGUAUAAGGAUCUAAGUAAAAUUUGUCGGUAUCUCGGUU-3’ for RNA. Fomivirsen, a fully phosphorothioated oligonucleotide therapeutic (sequence: 5′- GCGTTTGCTCTTCTTCTTGCG-3′), was obtained from MedChemExpress (Sollentuna, Sweden) in its sodium salt form. The sample was extensively desalted using repeated centrifugal microfiltration washing with 300 mM ammonium acetate, followed by several washing steps with MS grade water.

### Mass spectrometry

All mass analyses were performed on a prototype timsOmni platform (Bruker Switzerland AG, Fällanden, Switzerland) installed at the University of Geneva (Figure 2A). Samples solutions are injected at 5 µM or 10 µM oligonucleotide, at a flow rate of 2 µL/min. Parent ions for the MS² step are typically selected using the quadrupole mass filter upstream of the Omnitrap platform. The quadrupole mass filter RF driver has been modified extending isolation up to 4500 Th. The Omnitrap platform is a segmented linear ion trap consisting of 9 quadrupoles (Q1—Q9), defining three trapping regions for processing ions.^59^ All nine segments are driven by a pair of phase-coherent frequency-controlled antiphase rectangular waveforms applied at a fixed amplitude of 250V_0p_. The pressure established throughout all trapping sections is controlled dynamically using two fast pulse valves, each operated with a maximum repetition rate of 100 Hz.^59^ In Q2, ion isolation (for MS^n^ experiments) can be performed either using resolving DC signal components or using notched AC excitation signals. Resonance excitation CID (abbreviated _R_CID) is also carried out in Q2, using a short N_2_ gas pulse (the residence time of the collision gas is below 15 ms). Both _R_CID and thermalization following ion transfer to a neighboring segment can be accomplished within a single gas pulse. In Q5, the electron kinetic energy can be fine-tuned in the range 0—100 eV, with an energy spread of < 5eV.^60^ Finally, in Q8, laser windows were installed on either side of the vacuum housing accommodating the Omnitrap platform for coupling of both a UV and an IR laser in an orthogonal setup, unlike the coaxial arrangement tested previously.^61^ A fused silica window was installed on one side of the vacuum housing, providing optical access, for coupling with a UV-vis laser. A second BaF_2_ window was installed on the opposite side for coupling of an infrared CO_2_ laser.

**Figure 2:**
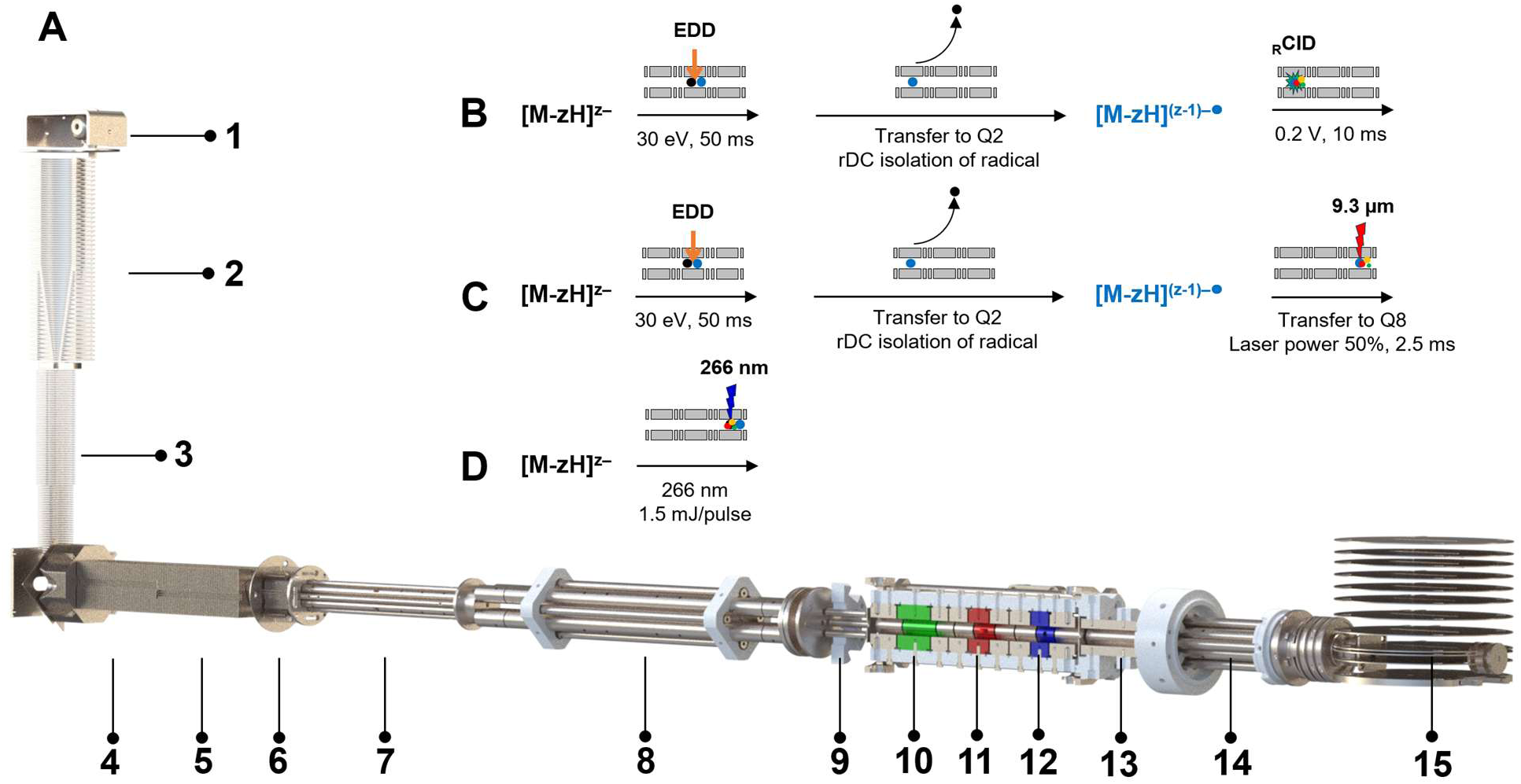
Schematic representation of the timsOmni mass spectrometer, highlighting the different processing steps explored for activating and dissociating ions. **(A)** The timsOmni MS platform consists of (1) a 18 cm long, 1 mm internal diameter resistive glass ion inlet receiving ions from an electrospray ionization source, (2) a RF ion funnel operated at 7 mbar, (3) a stacked-ring RF ion guide configured with an exit lens for declustering low charge state nucleic acids ions and performing collision induced unfolding at higher ion activation energies (4) an ion accumulation region, (5) a trapped ion mobility analyzer region (1.9 mbar), (6) a low pressure ion funnel enabling in-source CID, (7) a quadrupole ion guide for ion beam thermalization and with axial DC field for enhanced ion beam conditioning, (8) a quadrupole mass filter, (9) an RF hexapole ion guide. The Omnitrap platform consists of (10) section Q2 for ion selection based on resolving DC isolation and resonance excitation for _R_CID, (11) section Q5 for electron-based ion activation dissociation (ExD), with additional resolving DC and _R_CID capabilities for EXciD experiments and (12) section Q8 configured with optical access for laser irradiation. Each of sections Q2, Q5 and Q8 can be utilized for accumulating ions prior to activation. Processed ions are transferred by (13) a quadrupole ion guide Q10 into (14) a collision cell and ultimately sampled by (15) an orthogonal acceleration region equipped with DC ion optics for shaping the ion beam into a TOF mass analyzer. (**B**) **EDD-_R_CID MS**^3^: EDD in Q5 followed by transfer of the ions to Q2 where the radical ions are isolated using resolving DC. The radicals are then submitted to resonance CID in Q2 and sent to the TOF. (**C**) **EDD-IRMPD MS**^3^: is a variant where the radical ions formed by EDD are selected and sent to Q8 for infrared laser irradiation (IRMPD). (**D**) **UVPD**: the precursor ions are first accumulated in Q2 then sent to Q8 for UV laser irradiation.

We focused here on the activation of intact oligonucleotides in the negative ion mode and exploited several new combinations of ion activation methods available on the Omnitrap platform, some of which are schematically described in Figure 2B—D. For example, for top-down characterization of oligonucleotides using radical-based fragmentation, the precursor ion [M-zH]^z‒^ is selected and accumulated in Q5 for 100 ms in the presence of nitrogen gas. Typically, the accumulation period can be extended to several seconds or until the space charge limit of ∼50 M charges is reached. Radical ions are produced by EDD (irradiation with 30 eV electrons using typical irradiation times of 50—100 ms), the product ions are transferred to Q2 and the radical ions [M-zH]^(z-1)‒•^ are isolated by the application of a resolving DC (rDC) signal. The isolation process in Q2 involves a phase-coherent frequency jump of the rectangular waveforms and subsequent application of the rDC signals (±51 V) for parking ions at the tip of the stability diagram for a period of <1 ms. Any isolation step is applied 20 ms after a N_2_ gas pulse. An isolation width of ∼4 m/z is typically obtained in all experiments without ion losses. Alternatively, ions can be isolated in Q2 by applying AC frequency sweep excitation waveforms (20 ms) designed with a single frequency notch corresponding to the secular frequency of oscillation of the radical ion. The dissociation of [M-zH]^(z-1)‒•^ is then performed using either _R_CID in Q2 (Figure 2B), or the ions can be transferred to Q8 for laser irradiation (IRMPD, Figure 2C). The laser employed in this study is a Synrad V30i (Novanta, USA), operated at 9.3 µm wavelength, which is selected due to the enhanced absorption efficiency by nucleic acids.^62^ Typical irradiation times were between 1.5 and 7 ms at 50% power. In UVPD experiments (Figure 2D), the laser pulse frequency is synchronized with the processing cycle performed in the Omnitrap platform to define the number of laser pulses injected in section Q8. An electromechanical shutter is further employed to admit the laser beam into Q8. The UV laser (Continuum Powerlite 8010, Nd-YAG) operates at 355 nm to pump an OPO laser. Here we used the residual 532 nm of the pump laser and a frequency-doubling YAl_3_(BO_3_)_4_ crystal to generate a 266 nm beam (between 1.2 and 1.5 mJ/pulse), which is routed through Q8 using dielectric coated mirrors. In addition, the beam is shaped by a telescope built using a set of achromatic lenses. The laser beam has a diameter of 1.5 mm before entering section Q8, eliminating reflection off surfaces near or at the entrance and exit apertures (2.5 mm diameter) on the pole-electrodes.

### Data analysis

All raw spectra were processed using Bruker DataAnalysis 6.2 software ((Bruker, Bremen, Germany). Peak picking was achieved using the in-built Sum-peak algorithm with a S/N threshold, relative and absolute intensity all set to 0%. The resulting peak list is exported as a .csv file for further data treatment using open-source software tools.

For Fomivirsen, we used FAST-MS,^63^ which allows to incorporate chemical modifications to the phosphate backbone and define novel fragment ion types. The molecular properties were updated with the chemical formulas of the four sulfur-substituted nucleotides (A, T, G, C). Fragmentation lists were customized for both vibrational and radical-based experiments. Radical fragments are distinguished by a one-hydrogen loss relative to their closed-shell counterparts. The precursor ion was defined either as an intact closed-shell species for vibrational activation or as “intact-e” for radical-based processing workflows. Obtained results were exported in .xlsx format for further analysis. To eliminate poor quality ions and to minimize false-positive assignments, a two-tier filtering approach was applied to constitute the “assigned peaks” list presented in Table 1 (see results and discussion section). The first filter excluded ions with a quality factor below 0.2, while the second retained only those with a signal-to-noise ratio greater than 100. The theoretical masses of the fragments were calculated based on the chemical formula from the FAST- MS output.

**Table 1:** Summary of total number of peaks and product ions observed by six fragmentation methods for Fomivirsen.

|  | MS<sup>2</sup> |  |  |  | MS<sup>3</sup> |  |
| --- | --- | --- | --- | --- | --- | --- |
|  | RCID | IRMPD | UVPD | EDD | EDD-IRMPD | EDD-RCID |
| Total number of peaks | 531 | 472 | 572 | 632 | 199 | 244 |
| Number of assigned peaks | 249 | 211 | 189 | 222 | 65 | 109 |
| % assigned peaks | 46.9 | 44.7 | 33.0 | 35.1 | 32.7 | 44.7 |
| Among assigned peaks: |  |  |  |  |  |  |
| % 5' fragments (including *a*-B) | 49.4 | 42.2 | 49.7 | 64.9 | 60.0 | 70.6 |
| % 3' fragments | 20.1 | 19.9 | 29.6 | 31.1 | 33.8 | 28.4 |
| % other fragments + base losses | 30.5 | 37.9 | 20.6 | 4.1 | 6.2 | 0.9 |
| % radical fragments | 0.0 | 0.0 | 22.2 | 39.6 | 24.6 | 40.4 |
| Predominant product ion types | *a*-B, *a*, *y* | *a*-B, *a*, *y* | *a*, *w* | *a*, *d* | *a*, *a*<sup>•</sup>, *w*, *w*<sup>•</sup> | *a*, *a*<sup>•</sup>, *w*, *w*<sup>•</sup> |
| Sequence coverage | 100% | 100% | 100% | 100% | 100% | 100% |

For the analysis of 46-mer oligonucleotides, fragment assignments were performed using the Aom²S software.^64^ Spectral filters were set as follows: Intensity threshold of 0.001%, mass accuracy tolerance of 10 ppm with the condition that the monoisotopic peak is present, a minimum isotopic similarity of 80%, and a comparison mass zone ranging from -4 to 4. To detect radical ions, a variable group corresponding to hydrogen loss (H-1) was included. Assignment results were exported as Excel files and checked manually by comparing experimental isotopic distributions with theoretical predictions.

Theoretical fragment masses were calculated using the chemical formulas provided either by FAST- MS or Aom²S output files. Monoisotopic peak masses were determined based on the Bruker Isotope Pattern software, which served as the reference for calculating mass errors. This approach was applied consistently across all supplementary mass accuracy tables.

## RESULTS AND DISCUSSION

### EDD on DNA homo-hexamers

The short DNA hexamers dG_6_, dA_6_, dC_6_ and dT_6_ have been quite extensively studied in the past, using various activation techniques including EDD in FTICRMS^39, 65^ and UV irradiation in helium-filled quadrupole ion traps,^46, 66^ and thus constitute good benchmark systems in our preliminary testing of EDD in the timsOmni platform. The seminal FTICRMS work using a hollow cathode was conducted with electron kinetic energies in the range of 16 to 18 eV and electron irradiation times of 1 second.^36, 39, 65^ Later work by the Breuker group on longer RNA oligonucleotide used electron energies of 18 to 24 eV and irradiation times of 0.15 to 0.8 seconds.^37^ On the timsOmni, our optimal electron kinetic energy setting was 30 eV, and Figure 3 shows the radical ion yield resulting from electron detachment from the parent ions dB_6_^3-^, as well as the total fragment ion yields, as a function of the EDD irradiation time. The fragment ions observed here are identical to those that were observed previously on an FTICRMS. The EDD (MS²) mass spectra are shown in supporting information Figure S1. The propensity towards electron detachment (Figure 3A) also follows the previously observed EDD trend: dG_6_ > dT_6_ > dC_6_ > dA_6_. The main result is that, without optimization of the overlap between the ion cloud and the electron beam in these preliminary experiments, a high reaction yield is observed with irradiation times of the order of 100 ms, and the same products are observed as in FTICRMS. Therefore, despite different electron kinetic energy settings in Omnitrap and FTICR, the EDD effect is similar.

**Figure 3:**
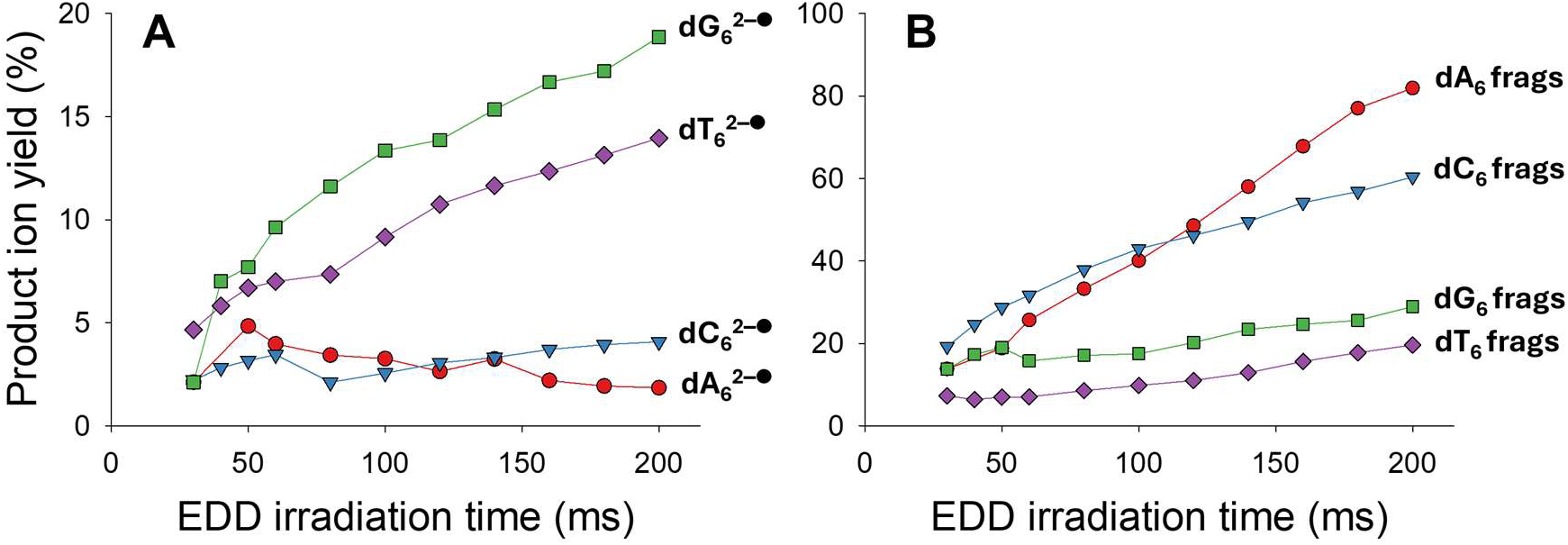
EDD on homo-hexamer DNAs. Product ion yields upon electron irradiation of dG_6_^3-^, dA_6_^3-^, dC_6_^3-^ and dT_6_^3-^, with an electron kinetic energy of 30 eV and as a function of the electron irradiation time, for two classes of product ions: A) products resulting from electron detachment (only single detachment was observed), and B) products resulting from covalent fragmentation.

### Fomivirsen therapeutic oligonucleotide: comparison of various activation techniques

Electrospraying Fomivirsen from 1mM NH_4_OAc produced high charge states (full scan MS spectrum in supporting information Figure S2). Here we compared a variety of options for ion activation on Fomivirsen 9‒ closed and open shell precursor ions (hence with about one charge per two nucleotides, which could be produced in high abundance. We compared four MS^2^ approaches on [M − 9H]^9−^ (vibrational activation with _R_CID and IRMPD—supporting information Figure S3, electronic activation with UVPD and EDD—Figure 4) and two MS^3^ approaches on [M − 10H]^10−^→[M − 10H]^9−•^ (EDD-_R_CID and EDD-IRMPD—Figure 5). Note that as the sequence is palindromic for the first and last 5 nucleotides, sometimes product ion identities cannot be deduced unambiguously as some fragment ions are isobaric. The isobaric palindromic pairs are *d* with *w*, *b* with *y*, *a* with *z*, and *c* with *x*. In such cases, the fastMS software returns a comment labeled “iso,” indicating isobaric, and it is up to the user to decide which ion to retain and which to discard. To resolve this, we confirm the identity of the first and last five nucleotides using fragment ion types that are not isobaric in the center of the sequence. Table 1 summarizes the types of product ion peaks observed in each method.

**Figure 4:**
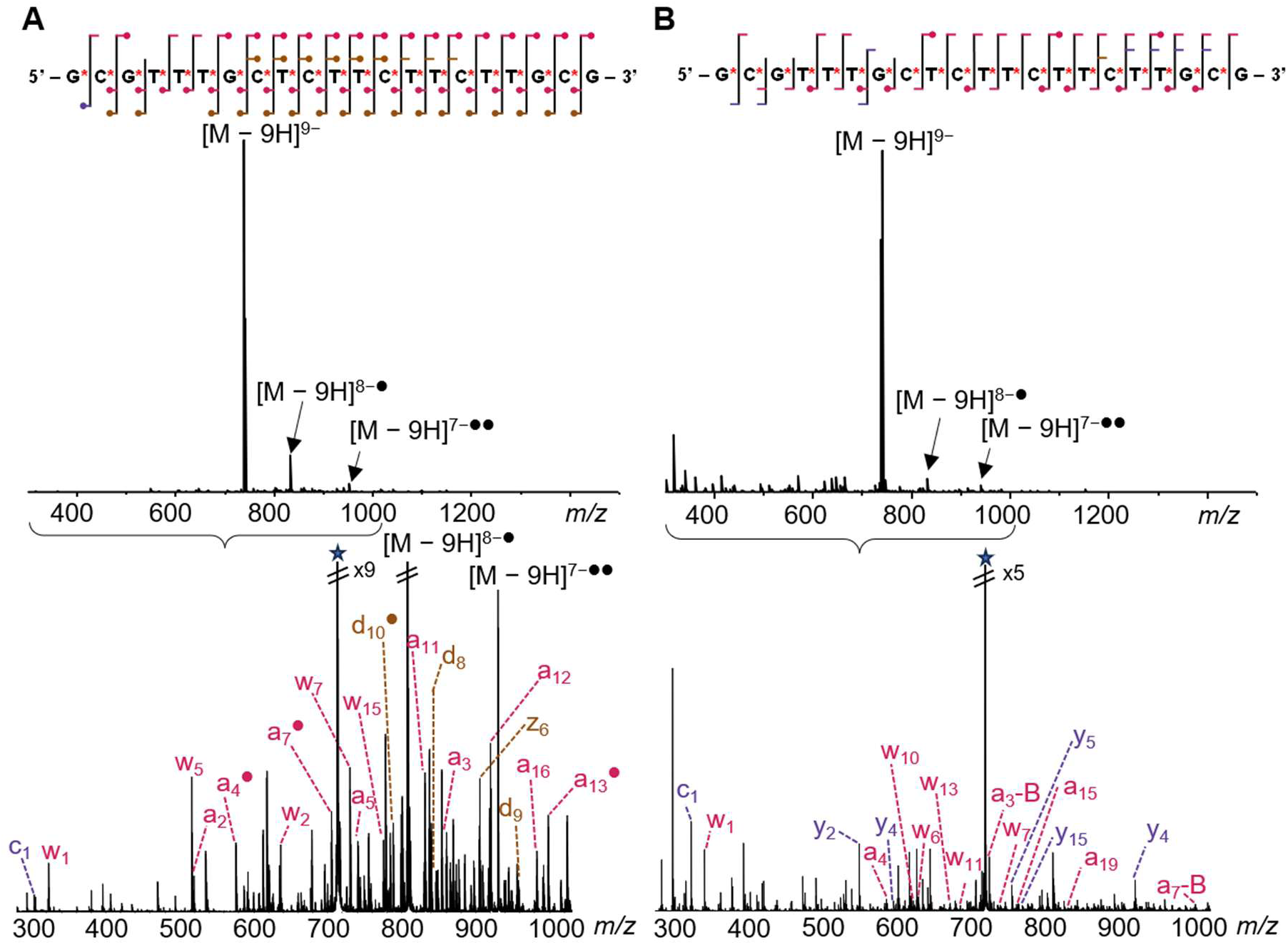
**(A) EDD** (28 eV, 55 ms) and **(B) UVPD** (266 nm, 10 Hz, 2 pulses, 1.5 mJ/pulse) **MS**^2^ spectra of Fomivirsen [M − 9H]^9−^, with sequence coverage map and main fragments annotated with the conventions of Figure 1. The star represents the parent ion. The fragment charge states were not written for clarity, and not all fragment families are represented on the sequence coverage map; see supporting information Tables S1 and S2 for the fragments serving for sequence coverage, and Table S7 for the full list of fragments. When the radical symbol is shown in the sequence coverage map, usually both the closed shell and the radical fragment are present.

**Figure 5:**
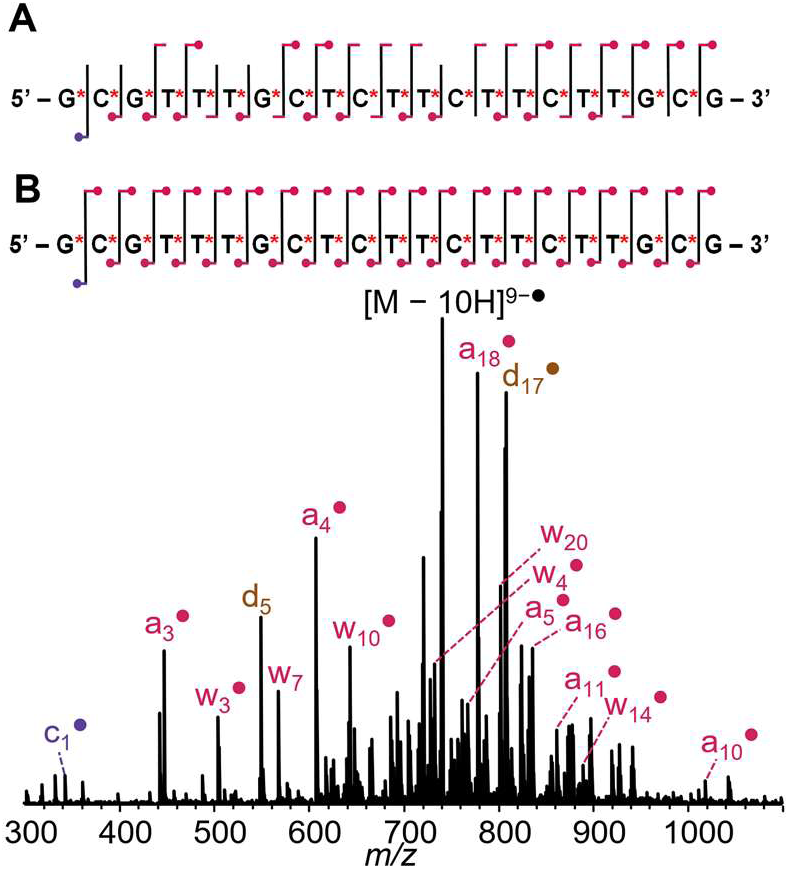
Sequence coverage map of **Fomivirsen** [M − 10H]^10−^→[M − 10H]^9−•^ using **(A) EDD-IRMPD MS**^3^ **and (B) EDD-_R_CID MS**^3^, for which an annotated spectrum is shown. See supporting information Tables S3 and S4 for the fragments serving for sequence coverage, and Table S7 for the full list of fragments.

All activation methods yielded 100% sequence coverage, but with different characteristics. In vibrational activation (_R_CID and IRMPD), *a*, *a*-B and *y* ions predominate and were sufficient to obtain full sequence coverage. The assigned peaks included all terminal backbone fragments, plus possible loss of one base. Unassigned peaks therefore include internal fragments and terminal fragments that would have lost more than one base. The mass spectra are very rich, with all eight fragment ion series detectable. Numerous fragments with base losses are also observed (30.5% of the assigned peaks), *y*-B ions being the most prominent after *a*-B. These observations are consistent with previous work on phosphorothioate oligonucleotides, which reported predominance of *a*-B and *w* fragment ions in vibrational activation modes,^67^ while a mixture of *d*, *w*, *c*, *y* and *x* ions was reported when using HCD and UVPD experiments.^49^ However, to the best of our knowledge, base loss fragments such as *y*-B and *w*-B have not been previously reported for fully phosphorothioate-modified oligonucleotide therapeutics.

EDD and UVPD (MS²) generated also very rich spectra, and the fraction of assigned peaks was lower than for vibrational activation MS². Yet full sequence coverage could be obtained by considering only *a* and *w* product ions for UVPD, and *a* and *d* product ions for EDD. Note that we carried out UVPD at 266 nm, a wavelength where single-photon electron photodetachment should occur, but here we used high laser energy per pulse and a focused beam to enhance the fragmentation yield. It is thus probable that we combine direct UVPD fragments, fragments coming from further dissociation of photodetachment products, and UVMPD (internal energy redistribution of multiple UV photons resulting in vibrational activation). Further work is needed to investigate the effect of the laser wavelength and energy on the balance of fragmentation pathways. EDD provided richer spectra, and also more electron detachment compared to our specific UVPD conditions. It also produced more radical fragments. The *a* and *w* series were the most prominent, followed by *d* and *z*, and all series were usually observed both in their closed shell and open shell forms (supporting information Figure S4).

Vibrational re-activation of the radical product ions formed by EDD produced overall fewer peaks, but despite the 10-fold drop in intensity due to the 10% radical ion yield in the EDD step, complete sequence coverage was obtained. IRMPD and _R_CID resulted in similar fragments, as expected. In particular, EDD- _R_CID MS^3^ presents several advantages compared to MS². The higher fraction of assigned peaks can be explained by the selective nature of the resonance collisional activation step targeting only the parent radical ion, in contrast to IRMPD where first-generation fragments can undergo further IR activation, resulting in secondary fragmentation and increasing the number of unassigned peaks. The fraction of radical ions is also particularly high, with complete complementary series of *a*•/*w* and *a*/*w*• ions in MS^3^. The irradiation conditions in IRMPD may require further optimization, but so far, EDD-_R_CID MS^3^ appears particularly promising. Ongoing work is devoted to extending the applicability of the Omnitrap ion activation network to oligonucleotides with other modified backbones, and establishing standardized methods needed for the detailed characterization of oligonucleotide therapeutics.

### EDD-based activation of 46-mer DNA and RNA

The next challenge is to examine longer DNA and RNA sequences (Figure 6). We first tested resonance _R_CID and IRMPD on closed-shell ions issued from electrospray. Both methods induce vibrational excitation and yield remarkably similar fragmentation patterns (fragment types, relative abundances, and sequence coverage; see Supplementary Table S8), therefore, only one is detailed in the main text. The observed fragment ions, *a*-B/*w* for DNA and *c*/*y* for RNA—are consistent with previous studies.^55–57^ The sequence coverage is incomplete at thymines for DNA (60% coverage), because thymines hamper the *a*-B/*w* fragmentation channel on their 3’ side. However, by inferring thymine presence from missing fragments and knowing its mass, the full sequence is resolved except for a single cleavage site. Owing to the high sensitivity of the timsOmni platform, acceptable sequence coverage is obtained with IRMPD, especially for RNA (91%). Improved results can be obtained, however, when combined with EDD.

**Figure 6:**
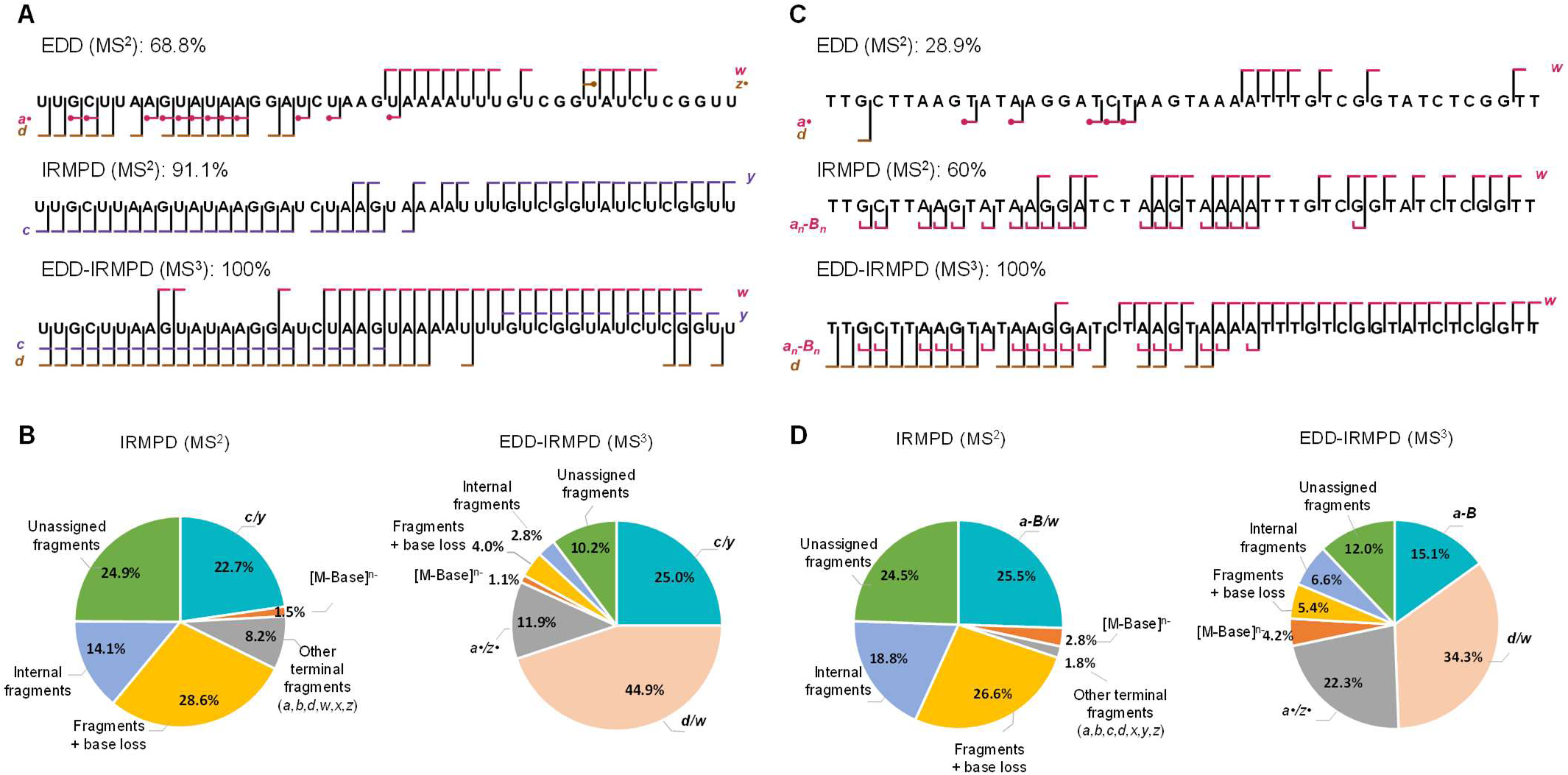
Comparison of 46-mer RNA. **(A) and DNA (C) sequence coverage** with EDD alone on [M- 8H]^8‒^, IRMPD alone on [M-8H]^8‒^, and EDD on [M-9H]^9‒^ followed by IRMPD on re-isolated [M-8H]^8‒•^. **(B,D): Fraction of peaks assigned to each fragment category**, for RNA and DNA respectively. The fragments indicated in bold are those used to map the sequence coverage. Unassigned fragments refer to ions that do not correspond to any terminal fragments, internal fragments, or base loss ions. Fragments within > 10 ppm were also considered as unassigned fragments. All fragment assignments are provided in supporting information Table S8. Note that due to the palindromic nature of the sequence, fragments corresponding to the first two positions are isobaric and cannot be separated by mass. Activation energies: EDD was performed using 30 eV electron kinetic energy and 50 ms irradiation time for both RNA and DNA. Vibrational activation was achieved using 6.6 ms IRMPD for RNA and 6 ms for DNA (9.3 µm, 50% power). For MS^3^ experiments, EDD was followed by IRMPD with 3 ms irradiation for RNA and 4.5 ms for DNA.

EDD MS² alone does not provide sufficiently high sequence coverage for the 46-mers, and the sequence coverage obtained for the 46-mer DNA (28.9%, see Figure 6C) is lower compared to the 46-mer RNA (68.8%, see Figure 6A). We note that we isolated a [M-9H]^9‒^ ion instead of higher charge states typically recommended,^37^ and thus any remaining intramolecular hydrogen bonds may hamper fragment ion formation or separation. The main product ions result from electron loss. However, upon re-activation of the [M-8H]^8‒•^ radical ions using IRMPD or ^R^CID, a rich product ion spectrum is produced, which allows to obtain full sequence coverage based on *d*/*w* and *c*/*y* ion series alone for RNA, and based on *d*/*w* and *a*- B/*w* for DNA. Notably, activation of open-shell DNA ions enables fragmentation on the 3’ side of thymines, which leads to full sequence coverage. *a*• and *z*• fragments were detected as well, but in low quantities and they are not essential to the sequence coverage.

EDD-IRMPD and EDD-_R_CID have the advantage of producing more sequence-informative fragments and fewer double fragments compared to vibrational activation alone (Figure 6B for RNA and 6D for DNA). We didn’t notice major differences in the distribution of fragments between EDD-IRMPD and EDD- _R_CID (supporting information Figure S5). In vibrational activation, only about a quarter of the product ion peaks (*c*/*y* ions for RNA, *a*–B/*w* ions for DNA) are sequence-informative. The others were either terminal fragments that sparingly confirm the sequence coverage, fragments that have lost a base (not necessarily adjacent to the cleavage site) or internal fragments. Under EDD-IRMPD and EDD-_R_CID conditions, additional informative *a*•/*w* and *d*/*z*• fragments are produced, resulting in complete sequence coverage.

These combined activation techniques also reduce internal fragmentation and non-informative base losses, simplifying spectral interpretation and increasing the proportion of sequence-informative fragments to 81.8% for RNA and 71.7% for DNA.

As noted above, *a*• and *z*• fragments were detected as well, but in low quantities. Furthermore, we observed that their abundance decreases when the IRMPD irradiation time is increased (see example(s) in Supporting Information Figure S6). We therefore wanted to determine the nature of the product ions when selecting *a*• and *z*• fragments as precursor ions. These experiments are challenging because of the low abundance of *a*• and *z*• fragments in 46-mers, but we managed to carry out both MS^3^ experiments on *a*• fragments produced by EDD on the 46-mer RNA (supporting information Figure S7), and an MS^4^ experiment on a *z*• fragment (Figure 7) or an *a*• fragment (supporting information Figure S10) re-isolated after EDD and _R_CID on the 46-mer DNA. The experiments showed that *a*_10_^3‒●^ from RNA dissociates into *d*_9_^3‒^, as previously postulated,^37^ but that the dissociation of *z*_6_^2‒●^ or *a*_8_^2‒●^ from DNA is more complicated and produces a variety of closed-shell ions. Yet overall, subsequent dissociation of *a*• and *z*• fragments upon vibrational activation explains the excess of closed-shell product ions compared to open-shell ones, even when starting from 100% re-isolated open-shell precursor ions.

**Figure 7:**
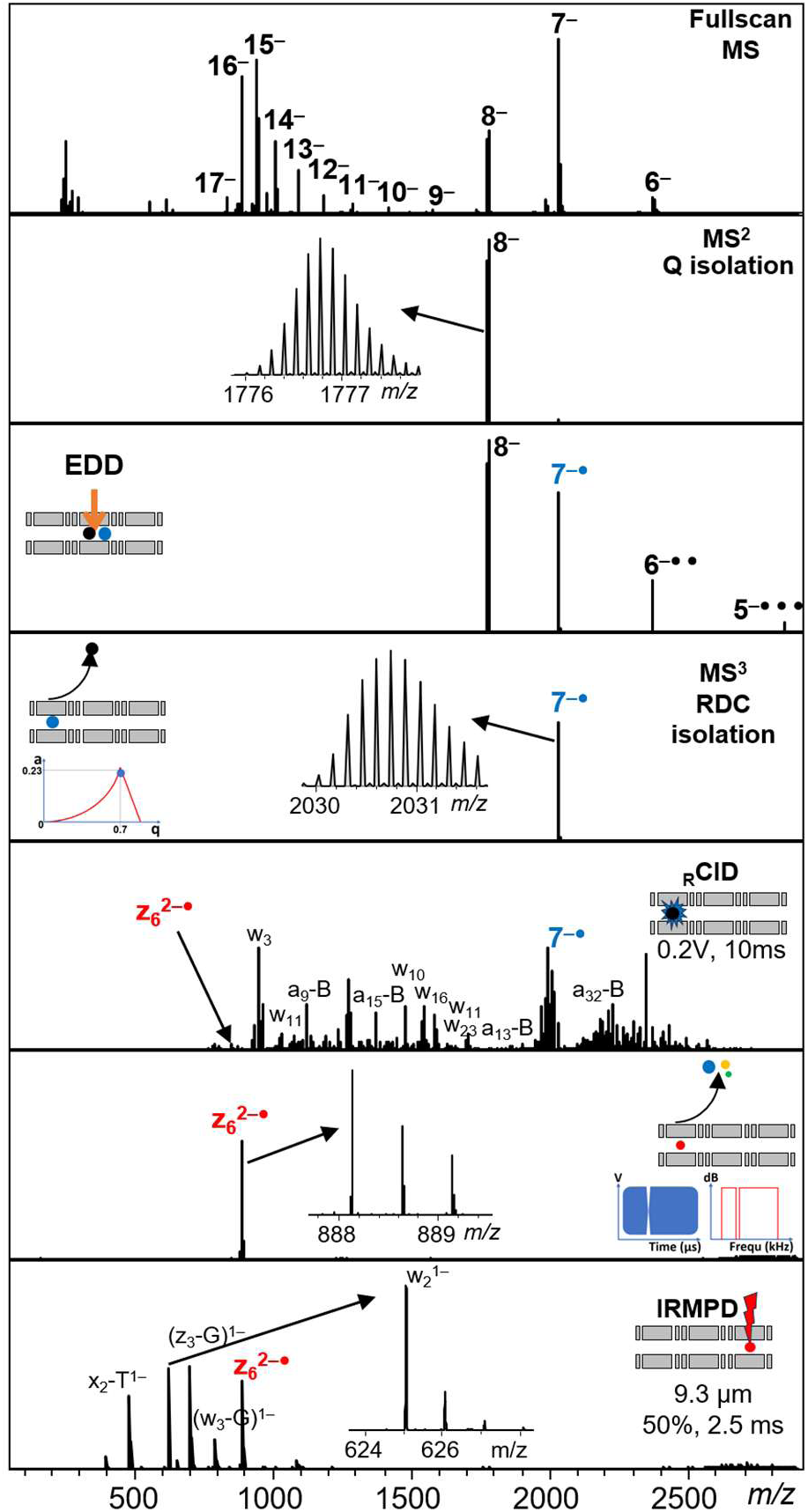
**Complete description of an MS**^4^ **experiment.** From the full scan MS spectrum, the charge state 8‒ (m/z 1776.9) is isolated using the quadrupole. A description Q1—Q9 DC gradient programming for each step is provided in supporting information Figure S8. The ions are accumulated in Q5 for 100 ms and submitted to EDD. 150 ms of irradiation with electron of kinetic energy 30 eV is used to produce radicals (the dependence of radical product ion yields on the irradiation time is shown in supporting information Figure S9). The radical 7^‒●^ (m/z 2030.7) is isolated in a MS^3^ stage using resolving DC by changing the drive frequency of the Omnitrap. The ions then undergo resonance CID during 10 ms (0.2 V). For the MS^4^ stage, the radical fragment z_6_^2‒●^, which is a very minor product in MS^3^, is re-isolated using a sweep of frequencies including a notch corresponding to its secular frequency. After transfer to section Q8, the radical fragment is irradiated for 2.5 ms (IRMPD, 9.3 µm, 50% power). The resulting fragments are a variety of closed shell ions. The total duration of the sequence is 350 ms, and the reported MS^4^ spectrum was recorded in 3 minutes.

## CONCLUSIONS

Our results highlight versatility of the Omnitrap platform for top-down characterization of intact, multiply charged oligonucleotide anions. EDD in the Omnitrap produces fragments similar to those obtained on other instrumental platforms. Both electron detachment and fragmentation occur, and the fraction of fragmentation products decreases with the oligonucleotide size, suggesting that subsequent fragmentation depends on intramolecular vibrational energy redistribution (IVR) following EDD. However, the EDD conditions in the Omnitrap platform are soft enough for radical fragments to survive, and therefore it becomes important that annotation software packages take into consideration the formation of these open-shell fragments. We also showed that upon vibrational re-activation or if sufficient energy is available through IVR, radical *a*• and *z*• fragments can further dissociate to form closed-shell ions.

EDD alone can still provide complete sequence coverage on a 21-mer phosphorothioate oligonucleotide therapeutic, but the direct EDD fragmentation efficiency decreases for longer oligonucleotides, and is lower for DNA than for RNA. Intentional vibrational re-activation of the parent ion radicals is thus required to increase the sequence coverage of EDD. On the Omnitrap platform, this can be carried out using MS^3^ workflows such as EDD-IRMPD and EDD-_R_CID, which result in better sequence coverage and less undesired secondary fragments. This is essential to characterize longer sequences. EDD- IRMPD is particularly fast to implement since no tuning or calibration of the rDC and ^R^CID steps is required, while EDD-^R^CID appears sufficiently soft, preserving radical fragments and eliminating secondary fragmentation processes.

We showed that the Omnitrap platform is amenable to combining different ion activation methods. Note that UVPD was only briefly discussed here, and further studies are required to explore ion dissociation pathways as a function of the laser wavelength and fluence, and gas pressure. The present work opens promising avenues in the field of oligonucleotide characterization, which is particularly timely for oligonucleotide therapeutics characterization, and is equally applicable to the characterization of post-transcriptional modifications on RNA. Excellent sensitivity and complete sequence coverage were readily achieved for the 46-mer DNA and RNA, despite the relatively low charge states selected here (0.2 charges per phosphate). Future efforts will focus on extending this capability to larger sequences and studying the influence of precursor charge state on the sequence coverage in a more systematic manner. Besides, prospects for native top-down characterization of nucleic structures are currently being explored.

## SUPPORTING INFORMATION

Supplementary figures as described in the text include additional mass spectra, sequence coverage maps, and supplementary MS^3^ and MS^4^ experiments. Tables of fragments annotated on the Fomivirsen sequence coverage maps (Tables S1 to S6) (PDF).

Supplementary Table S7: full fragment lists for Fomivirsen (XLSX). Supplementary Table S8: full fragment lists for the 46-mer DNA and RNA (XLSX).

## ACKNOWLEDGEMENTS

This work was supported by the Swiss National Science Foundation (SNF R’Equip grant 220390 to VG), the Boninchi foundation, the Ernest and Lucie Schmidheiny foundation, the Société Académique de Genève, the University of Geneva, and by the Swiss State Secretariat for Education, Research and Innovation (SERI) for participation to the Marie Skłodowska-Curie Action doctoral network MobiliTraIN (grant number 101119562).

## CONLFICT OF INTEREST DISCLOSURE

The authors declare the following possible conflict of interest(s): A. Smyrnakis, J.-F. Greisch and D. Papanastasiou are employees of Bruker, which develops and manufactures advanced mass spectrometry platforms for life science applications. This article includes data and findings related to Bruker’s new timsOmni product.

