## Supplemental Figures S1-S10, supplemental Tables S1-S6 for "Activation of oligonucleotide polyanions using collisions, electrons and photons in a timsOmni platform"

SUPPORTING INFORMATION

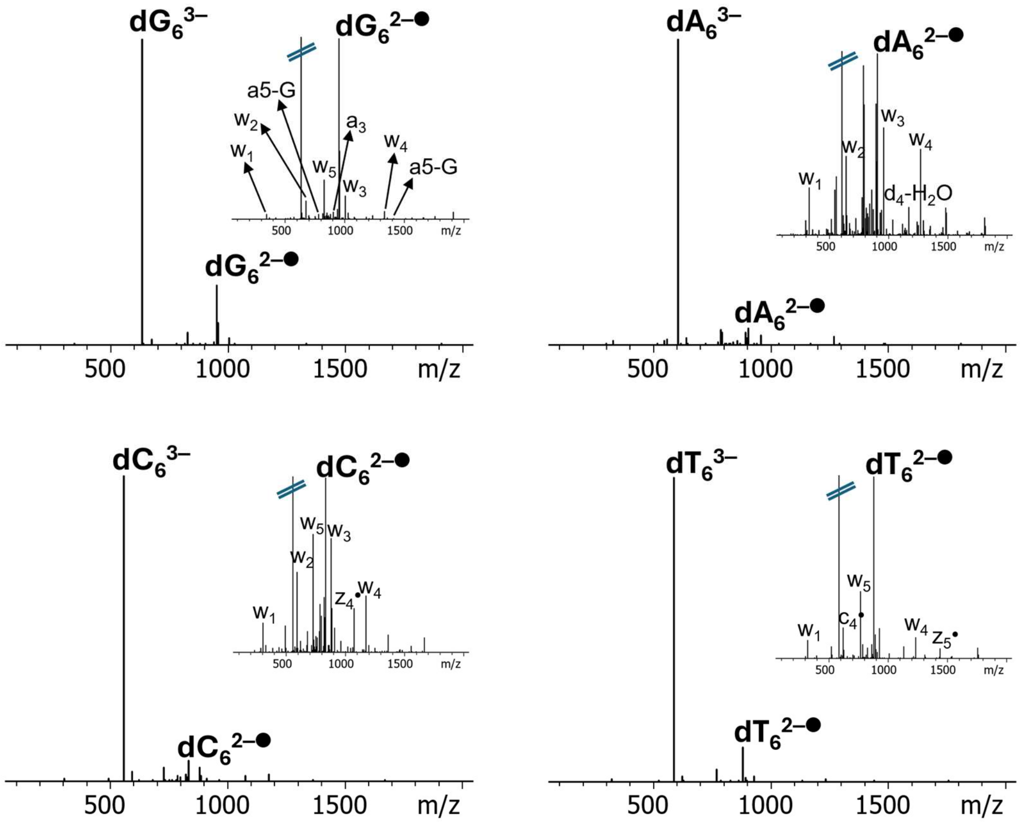


**Figure S1:** Product ion spectra resulting from EDD on DNA homo-hexamers $dG_6^{3-}$, $dA_6^{3-}$, $dC_6^{3-}$ and $dT_6^{3-}$.

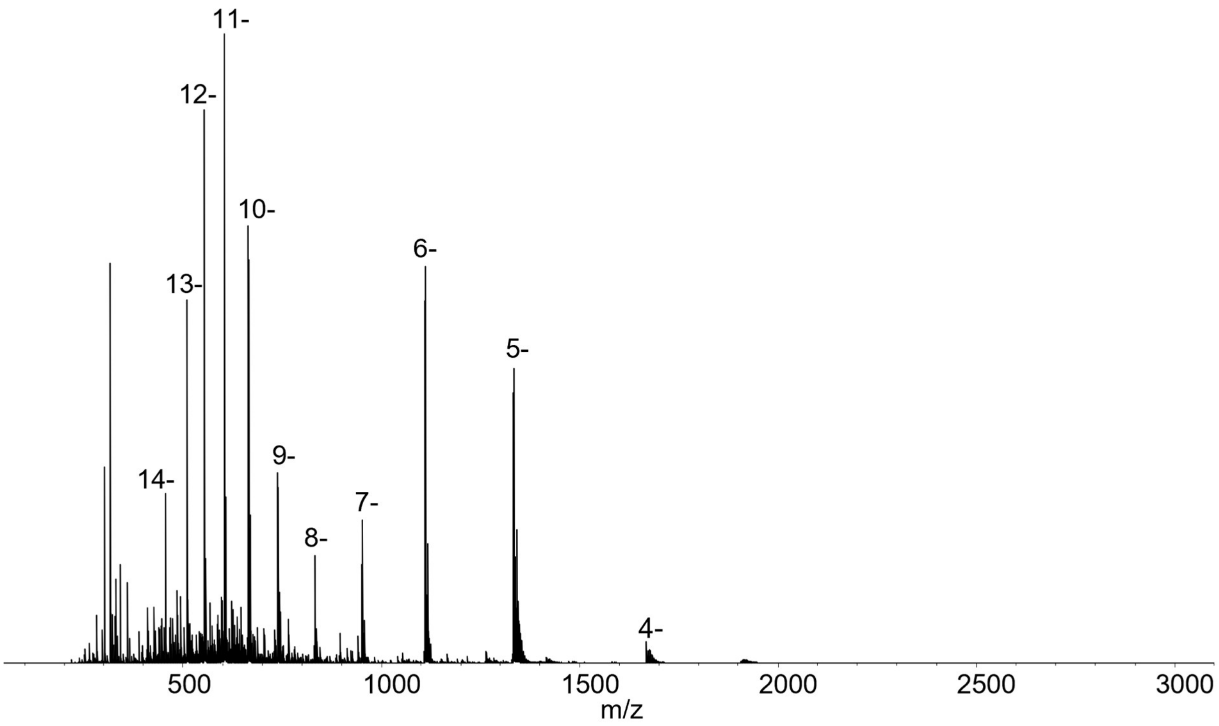


**Figure S2:** Full scan MS spectrum of Fomivirsen sprayed from 1 mM aqueous $NH_4OAc$, showing the initial charge state distribution.

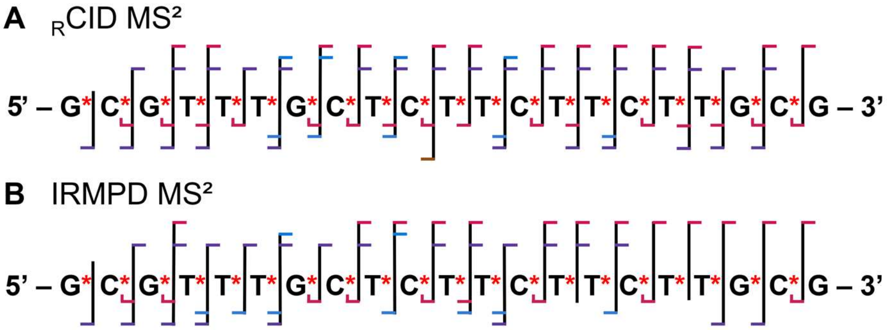


**Figure S3:** sequence coverage map of (**A**) $_{R}$CID MS² and (**B**) IRMPD MS² of Fomivirsen, $[M - 9H]^{9-}$, annotated using the nomenclature shown in Figure 1. $_{R}$CID amplitude and IR irradiation time were optimized to achieve informative fragmentation patterns while preserving some precursor ion intensity (on the order of ~≥50%). See Tables S5 and S6 for peak assignment with ppm errors. Only a subset of terminal fragment ions with high S/N are marked in the sequence coverage map. All other high/low intensity fragment ions of other charge states conforming the same positions are attached in supplementary Table S7.

**Table S1:** Peak list table on the fragments annotated on the sequence coverage map for Fomivirsen and that suffice to provide full sequence coverage, from the **EDD MS²** spectrum on $[M - 9H]^{9-}$. Full FAST MS output of fragment ion peaks of Fomivrisen $[M - 9H]^{9-}$ is provided in supplementary Table S7 (separate Excel file).

| nt | Position (5') | Peak assignment | Charge state (z) | Observed m/z | Theoretical m/z | Mass error (ppm) | nt | Position (3') | Peak assignment | Charge state (z) | Observed m/z | Theoretical m/z | Mass error (ppm) |
|---|---|---|---|---|---|---|---|---|---|---|---|---|---|
| **G** | 1 | c1 | 1- | 343.0139 | 343.0145 | -1.75 | **G** | 21 | | | | | |
| **C** | 2 | a2 | 1- | 553.1038 | 553.1024 | 2.53 | **C** | 20 | w20 | 7- | 917.3538 | 917.3502 | 3.92 |
| **G** | 3 | a3 | 1- | 898.132 | 898.1321 | -0.11 | **G** | 19 | w19• | 7- | 873.6311 | 873.6314 | -0.34 |
| **T** | 4 | a4• | 2- | 608.072 | 608.0701 | 3.12 | **T** | 18 | | | | | |
| **T** | 5 | a5• | 2- | 768.084 | 768.0817 | 2.99 | **T** | 17 | w17 | 6- | 908.7349 | 908.7303 | 5.06 |
| **T** | 6 | a6• | 2- | 928.0959 | 928.0933 | 2.80 | **T** | 16 | w16 | 6- | 855.3962 | 855.3931 | 3.62 |
| **G** | 7 | a7• | 3- | 733.4066 | 733.403 | 4.91 | **G** | 15 | w15 | 6- | 802.0596 | 802.0559 | 4.61 |
| **C** | 8 | a8• | 2- | 1253.1228 | 1253.1199 | 2.31 | **C** | 14 | w14 | 5- | 893.6653 | 893.6627 | 2.90 |
| **T** | 9 | d9 | 3- | 980.0766 | 980.0725 | 4.18 | **T** | 13 | z13 | 4- | 1012.5888 | 1012.5858 | 2.96 |
| **C** | 10 | a10• | 3- | 1043.4314 | 1043.4264 | 4.79 | **C** | 12 | w12 | 5- | 768.6559 | 768.6533 | 3.38 |
| **T** | 11 | a11 | 4- | 862.5785 | 862.5757 | 3.25 | **T** | 11 | z11 | 4- | 856.328 | 856.3241 | 4.55 |
| **T** | 12 | a12 | 4- | 942.5843 | 942.5815 | 2.97 | **T** | 10 | w10 | 4- | 804.8104 | 804.8068 | 4.47 |
| **C** | 13 | a13• | 4- | 1018.591 | 1018.5854 | 5.50 | **C** | 9 | z9 | 3- | 928.7554 | 928.7524 | 3.23 |
| **T** | 14 | a14 | 5- | 878.8761 | 878.8731 | 3.41 | **T** | 8 | w8 | 4- | 648.5483 | 648.5451 | 4.93 |
| **T** | 15 | a15• | 6- | 785.3981 | 785.3956 | 3.18 | **T** | 7 | w7 | 3- | 758.3904 | 758.3882 | 2.90 |
| **C** | 16 | a16 | 5- | 1003.6876 | 1003.6809 | 6.68 | **C** | 6 | z6 | 2- | 921.093 | 921.0973 | -4.66 |
| **T** | 17 | a17 | 6- | 889.7414 | 889.738 | 3.82 | **T** | 5 | w5 | 3- | 550.0416 | 550.0393 | 4.18 |
| **T** | 18 | a18• | 7- | 808.2105 | 808.2063 | 5.20 | **T** | 4 | w4• | 1- | 665.0487 | 665.0471 | 2.41 |
| **G** | 19 | a19 | 7- | 857.5008 | 857.4962 | 5.36 | **G** | 3 | w3 | 1- | 1012.0884 | 1012.0861 | 2.27 |
| **C** | 20 | a20 | 7- | 901.075 | 901.071 | 4.44 | **C** | 2 | w2 | 1- | 667.0579 | 667.0564 | 2.25 |
| **G** | 21 | | | | | | **G** | 1 | w1 | 1- | 362.0338 | 362.0329 | 2.49 |

**Table S2:** Peak list table on the fragments annotated on the sequence coverage map for Fomivirsen and that suffice to provide full sequence coverage, from the **UVPD MS²** spectrum on $[M - 9H]^{9-}$. Full FAST MS output of fragment ion peaks of Fomivrisen $[M - 9H]^{9-}$ is provided in supplementary Table S7 (separate Excel file).

| nt | Position (5') | Peak assignment | Charge state (z) | Observed m/z | Theoretical m/z | Mass error (ppm) | nt | Position (3') | Peak assignment | Charge state (z) | Observed m/z | Theoretical m/z | Mass error (ppm) |
|---|---|---|---|---|---|---|---|---|---|---|---|---|---|
| **G** | 1 | c1 | 1- | 344.0224 | 344.0224 | 0.00 | **G** | 21 | | | | | |
| **C** | 2 | c2 | 1- | 649.0467 | 649.0459 | 1.23 | **C** | 20 | w20 | 7- | 917.3509 | 917.3502 | 0.76 |
| **G** | 3 | a3 | 1- | 898.1318 | 898.1321 | -0.33 | **G** | 19 | | | | | |
| **T** | 4 | a4 | 2- | 608.5738 | 608.574 | -0.33 | **T** | 18 | | | | | |
| **T** | 5 | a5 | 2- | 768.5858 | 768.5856 | 0.26 | **T** | 17 | w17 | 6- | 908.7327 | 908.7303 | 2.64 |
| **T** | 6 | c6 | 3- | 650.7127 | 650.7102 | 3.84 | **T** | 16 | w16 | 6- | 855.3972 | 855.3931 | 4.79 |
| **G** | 7 | a7• | 3- | 733.4042 | 733.403 | 1.64 | **G** | 15 | y15 | 6- | 786.0649 | 786.0654 | -0.64 |
| **C** | 8 | a8 | 3- | 835.414 | 835.4134 | 0.72 | **C** | 14 | | | | | |
| **T** | 9 | | | | | | **T** | 13 | w13 | 6- | 693.7155 | 693.7137 | 2.59 |
| **C** | 10 | a10• | 3- | 1043.4345 | 1043.4264 | 7.76 | **C** | 12 | w12 | 5- | 768.6562 | 768.6533 | 3.77 |
| **T** | 11 | a11 | 4- | 862.5785 | 862.5757 | 3.25 | **T** | 11 | w11 | 5- | 707.6505 | 707.6486 | 2.68 |
| **T** | 12 | | | | | | **T** | 10 | w10 | 5- | 643.6472 | 643.644 | 4.97 |
| **C** | 13 | a13 | 5- | 814.8706 | 814.8684 | 2.70 | **C** | 9 | w9 | 4- | 724.805 | 724.801 | 5.52 |
| **T** | 14 | a14 | 5- | 878.8751 | 878.8731 | 2.28 | **T** | 8 | w8 | 4- | 648.5481 | 648.5451 | 4.63 |
| **T** | 15 | a15 | 6- | 785.5573 | 785.5635 | -7.89 | **T** | 7 | w7 | 3- | 758.3905 | 758.3882 | 3.03 |
| **C** | 16 | a16 | 6- | 836.409 | 836.4008 | 9.80 | **C** | 6 | w6 | 3- | 651.7151 | 651.7138 | 1.99 |
| **T** | 17 | a17 | 6- | 889.7408 | 889.738 | 3.15 | **T** | 5 | y5 | 2- | 777.5927 | 777.5908 | 2.44 |
| **T** | 18 | a18• | 7- | 808.2105 | 808.2063 | 5.20 | **T** | 4 | y4 | 2- | 617.5807 | 617.5793 | 2.27 |
| **G** | 19 | a19 | 7- | 857.5001 | 857.4962 | 4.55 | **G** | 3 | y3 | 1- | 916.1433 | 916.1426 | 0.76 |
| **C** | 20 | | | | | | **C** | 2 | y2 | 1- | 571.1135 | 571.113 | 0.88 |
| **G** | 21 | | | | | | **G** | 1 | w1 | 1- | 362.0329 | 362.0329 | 0.00 |

**Table S3:** Peak list table on the fragments annotated on the sequence coverage map for Fomivirsen and that suffice to provide full sequence coverage, from the **EDD-IRMPD MS$^3$** spectrum on $[M-9H]^{9-\bullet}$. Full FAST MS output of fragment ion peaks of Fomivrisen $[M-9H]^{9-\bullet}$ is provided in supplementary Table S7 (separate Excel file).

| nt | Position (5') | Peak assignment | Charge state (z) | Observed m/z | Theoretical m/z | Mass error (ppm) | nt | Position (3') | Peak assignment | Charge state (z) | Observed m/z | Theoretical m/z | Mass error (ppm) |
|---|---|---|---|---|---|---|---|---|---|---|---|---|---|
| **G** | 1 | c1• | 1- | 343.0169 | 343.0145 | 7.00 | **G** | 21 | | | | | |
| **C** | 2 | a2• | 1- | 552.096 | 552.0946 | 2.54 | **C** | 20 | | | | | |
| **G** | 3 | a3• | 1- | 897.1268 | 897.1242 | 2.90 | **G** | 19 | | | | | |
| **T** | 4 | a4• | 2- | 608.0742 | 608.0701 | 6.74 | **T** | 18 | w18 | 8- | 721.2998 | 721.2988 | 1.39 |
| **T** | 5 | a5 | 2- | 768.5872 | 768.5856 | 2.08 | **T** | 17 | w17• | 7- | 778.7744 | 778.7678 | 8.47 |
| **T** | 6 | a6• | 3- | 618.3945 | 618.3931 | 2.26 | **T** | 16 | | | | | |
| **G** | 7 | a7 | 3- | 733.7406 | 733.7389 | 2.32 | **G** | 15 | | | | | |
| **C** | 8 | a8• | 4- | 626.058 | 626.0563 | 2.72 | **C** | 14 | w14• | 6- | 744.3881 | 744.383 | 6.85 |
| **T** | 9 | a9• | 4- | 706.0647 | 706.0621 | 3.68 | **T** | 13 | w13• | 6- | 693.549 | 693.5458 | 4.61 |
| **C** | 10 | a10 | 4- | 1043.4315 | 1043.4264 | 4.89 | **C** | 12 | w12 | 5- | 768.6566 | 768.6533 | 4.29 |
| **T** | 11 | a11• | 4- | 862.3276 | 862.3237 | 4.52 | **T** | 11 | w11 | 5- | 707.6548 | 707.6486 | 8.76 |
| **T** | 12 | a12• | 5- | 753.6672 | 753.6622 | 6.63 | **T** | 10 | w10 | 5- | 643.6498 | 643.644 | 9.01 |
| **C** | 13 | | | | | | **C** | 9 | | | | | |
| **T** | 14 | a14• | 6- | 732.0628 | 732.0584 | 6.01 | **T** | 8 | w8 | 4- | 648.5502 | 648.5451 | 7.86 |
| **T** | 15 | a15• | 6- | 785.4004 | 785.3956 | 6.11 | **T** | 7 | w7 | 4- | 568.5434 | 568.5393 | 7.21 |
| **C** | 16 | a16 | 7- | 716.7726 | 716.7711 | 2.09 | **C** | 6 | w6• | 3- | 651.3795 | 651.3779 | 2.46 |
| **T** | 17 | a17• | 7- | 762.3469 | 762.3447 | 2.89 | **T** | 5 | w5 | 2- | 825.5655 | 825.5626 | 3.51 |
| **T** | 18 | a18 | 7- | 808.2106 | 808.2063 | 5.32 | **T** | 4 | w4• | 2- | 665.0492 | 665.0471 | 3.16 |
| **G** | 19 | | | | | | **G** | 3 | w3• | 2- | 505.0374 | 505.0355 | 3.76 |
| **C** | 20 | | | | | | **C** | 2 | w2• | 1- | 666.054 | 666.0486 | 8.11 |
| **G** | 21 | | | | | | **G** | 1 | w1• | 1- | 361.0221 | 361.0251 | -8.31 |

**Table S4:** Peak list table on the fragments annotated on the sequence coverage map for Fomivirsen and that suffice to provide full sequence coverage, from the **EDD-$_{R}$CID MS$^3$** spectrum on $[M - 9H]^{9-\bullet}$. Full FAST MS output of fragment ion peaks of Fomivrisen $[M - 9H]^{9-\bullet}$ is provided in supplementary Table S7 (separate Excel file).

| nt | Position (5') | Peak assignment | Charge state (z) | Observed m/z | Theoretical m/z | Mass error (ppm) | nt | Position (3') | Peak assignment | Charge state (z) | Observed m/z | Theoretical m/z | Mass error (ppm) |
|---|---|---|---|---|---|---|---|---|---|---|---|---|---|
| **G** | 1 | c1• | 1- | 343.0154 | 343.0145 | 2.62 | **G** | 21 | | | | | |
| **C** | 2 | a2• | 1- | 552.096 | 552.0946 | 2.54 | **C** | 20 | w20 | 8- | 802.561 | 802.5555 | 6.85 |
| **G** | 3 | a3• | 2- | 448.0614 | 448.0585 | 6.47 | **G** | 19 | w19• | 8- | 764.3053 | 764.3016 | 4.84 |
| **T** | 4 | a4• | 2- | 608.0742 | 608.0699 | 7.07 | **T** | 18 | w18 | 8- | 721.3019 | 721.2988 | 4.30 |
| **T** | 5 | a5• | 2- | 768.081 | 768.0817 | -0.91 | **T** | 17 | w17• | 7- | 778.7738 | 778.7678 | 7.70 |
| **T** | 6 | a6• | 3- | 618.3945 | 618.3931 | 2.26 | **T** | 16 | w16• | 7- | 733.0556 | 733.0502 | 7.37 |
| **G** | 7 | a7• | 3- | 733.407 | 733.403 | 5.45 | **G** | 15 | w15• | 7- | 687.188 | 687.1886 | -0.87 |
| **C** | 8 | a8• | 4- | 626.0589 | 626.0563 | 4.15 | **C** | 14 | w14• | 5- | 893.4615 | 893.4611 | 0.45 |
| **T** | 9 | a9• | 4- | 706.0645 | 706.0621 | 3.40 | **T** | 13 | w13• | 6- | 693.549 | 693.5458 | 4.61 |
| **C** | 10 | a10• | 3- | 1043.4334 | 1043.4264 | 6.71 | **C** | 12 | w12• | 5- | 768.4589 | 768.4517 | 9.37 |
| **T** | 11 | a11• | 4- | 862.3253 | 862.3237 | 1.86 | **T** | 11 | w11• | 5- | 707.4423 | 707.447 | -6.64 |
| **T** | 12 | a12• | 5- | 753.6673 | 753.6622 | 6.77 | **T** | 10 | w10• | 5- | 643.4409 | 643.4424 | -2.33 |
| **C** | 13 | a13 | 5- | 814.8735 | 814.8684 | 6.26 | **C** | 9 | w9 | 4- | 724.808 | 724.801 | 9.66 |
| **T** | 14 | a14• | 6- | 732.0603 | 732.0584 | 2.60 | **T** | 8 | w8 | 4- | 648.5481 | 648.5451 | 4.63 |
| **T** | 15 | a15• | 6- | 785.3941 | 785.3956 | -1.91 | **T** | 7 | w7 | 4- | 568.5422 | 568.5393 | 5.10 |
| **C** | 16 | a16• | 6- | 836.2386 | 836.2328 | 6.94 | **C** | 6 | w6• | 3- | 651.3804 | 651.3779 | 3.84 |
| **T** | 17 | a17• | 7- | 762.3475 | 762.3447 | 3.67 | **T** | 5 | w5 | 2- | 825.5644 | 825.5626 | 2.18 |
| **T** | 18 | a18• | 7- | 808.21 | 808.2063 | 4.58 | **T** | 4 | w4• | 2- | 665.052 | 665.0471 | 7.37 |
| **G** | 19 | a19• | 8- | 750.0509 | 750.0573 | -8.53 | **G** | 3 | w3 | 2- | 505.036 | 505.0355 | 0.99 |
| **C** | 20 | a20• | 8- | 788.1895 | 788.1852 | 5.46 | **C** | 2 | w2• | 1- | 666.0531 | 666.0486 | 6.76 |
| **G** | 21 | | | | | | **G** | 1 | w1• | 1- | 361.0221 | 361.0251 | -8.31 |

**Table S5:** Peak list table on the fragments annotated on the sequence coverage map for Fomivirsen and that suffice to provide full sequence coverage, from the $_{\mathbf{R}}$**CID MS**$^{2}$ spectrum on $[M-9H]^{9-}$. Full FAST MS output of fragment ion peaks of Fomivrisen $[M-9H]^{9}$ is provided in supplementary Table S7 (separate Excel file).

| nt | Position (5') | Peak assignment | Charge state (z) | Observed m/z | Theoretical m/z | Mass error (ppm) | nt | Position (3') | Peak assignment | Charge state (z) | Observed m/z | Theoretical m/z | Mass error (ppm) |
|---|---|---|---|---|---|---|---|---|---|---|---|---|---|
| **G** | 1- | c1 | 1- | 344.0228 | 344.0224 | 0.12 | **G** | 21 | | | | | |
| **C** | 2 | a2-C | 1- | 442.0598 | 442.0591 | 0.16 | **C** | 20 | y20 | 8- | 790.5602 | 790.5625 | -0.29 |
| **G** | 3 | a3-G | 1- | 747.085 | 747.0827 | 0.31 | **G** | 19 | y19 | 8- | 752.5641 | 752.5599 | 0.56 |
| **T** | 4 | c4 | 2- | 656.5482 | 656.5457 | 0.38 | **T** | 18 | y18 | 7- | 810.7843 | 810.7792 | 0.63 |
| **T** | 5 | b5 | 2- | 777.5939 | 777.5908 | 0.40 | **T** | 17 | y17 | 7- | 765.0583 | 765.0616 | -0.43 |
| **T** | 6 | c6 | 3- | 650.7138 | 650.7102 | 0.55 | **T** | 16 | y16 | 6- | 839.4066 | 839.4026 | 0.48 |
| **G** | 7 | a7-G | 2- | 1025.5921 | 1025.5873 | 0.47 | **G** | 15 | y15 | 6- | 786.0696 | 786.0654 | 0.53 |
| **C** | 8 | a8-C | 3- | 798.4022 | 798.399 | 0.40 | **C** | 14 | w14 | 6- | 744.5568 | 744.551 | 0.78 |
| **T** | 9 | b9 | 4- | 710.8206 | 710.8167 | 0.55 | **T** | 13 | w13 | 6- | 693.7173 | 693.7137 | 0.52 |
| **C** | 10 | a10-C | 4- | 754.8124 | 754.8091 | 0.44 | **C** | 12 | y12 | 5- | 749.4681 | 749.4646 | 0.47 |
| **T** | 11 | a11 | 4- | 862.5798 | 862.5757 | 0.48 | **T** | 11 | w11 | 5- | 707.6526 | 707.6486 | 0.57 |
| **T** | 12 | b12 | 5- | 757.4699 | 757.4658 | 0.54 | **T** | 10 | y10 | 4- | 780.8248 | 780.8209 | 0.50 |
| **C** | 13 | a13-C | 5- | 792.6643 | 792.6598 | 0.57 | **C** | 9 | y9 | 4- | 700.8184 | 700.8151 | 0.47 |
| **T** | 14 | a14 | 5- | 878.8772 | 878.8731 | 0.47 | **T** | 8 | y8 | 3- | 833.0823 | 833.0814 | 0.11 |
| **T** | 15 | b15 | 6- | 788.5686 | 788.5653 | 0.42 | **T** | 7 | w7 | 4- | 568.5428 | 568.5393 | 0.62 |
| **C** | 16 | a16 | 6- | 836.4138 | 836.4008 | 1.55 | **C** | 6 | w6 | 4- | 488.5367 | 488.5335 | 0.66 |
| **T** | 17 | a17 | 6- | 889.7431 | 889.738 | 0.57 | **T** | 5 | w5 | 3- | 550.0422 | 550.0393 | -0.53 |
| **T** | 18 | c18 | 7- | 821.9151 | 821.9125 | 0.32 | **T** | 4 | y4 | 2- | 617.5822 | 617.5793 | 0.47 |
| **G** | 19 | c19 | 8- | 762.1805 | 762.1762 | 0.56 | **G** | 3 | y3 | 2- | 457.5693 | 457.5677 | 0.35 |
| **C** | 20 | a20-C | 8- | 774.4526 | 774.4308 | 2.81 | **C** | 2 | w2 | 1- | 667.0582 | 667.0564 | 0.27 |
| **G** | 21 | | | | | | **G** | 1 | w1 | 1- | 362.0337 | 362.0329 | 0.22 |

**Table S6:** Peak list table on the fragments annotated on the sequence coverage map for Fomivirsen and that suffice to provide full sequence coverage, from the **IRMPD MS²** spectrum on $[M-9H]^{9-}$. Full FAST MS output of fragment ion peaks of Fomivrisen $[M-9H]^{9-}$ is provided in supplementary Table S7 (separate Excel file).

| nt | Position (5') | Peak assignment | Charge state (z) | Observed m/z | Theoretical m/z | Mass error (ppm) | nt | Position (3') | Peak assignment | Charge state (z) | Observed m/z | Theoretical m/z | Mass error (ppm) |
|---|---|---|---|---|---|---|---|---|---|---|---|---|---|
| **G** | 1 | c1 | 1- | 344.0233 | 344.0224 | 0.26 | **G** | 21 | | | | | |
| **C** | 2 | a2-C | 1- | 442.0599 | 442.0591 | 0.18 | **C** | 20 | | | | | |
| **G** | 3 | a3-G | 1- | 747.0845 | 747.0827 | 0.24 | **G** | 19 | y19 | 8- | 752.4386 | 752.4346 | 0.53 |
| **T** | 4 | c4 | 2- | 656.5488 | 656.5457 | 0.47 | **T** | 18 | y18 | 7- | 810.7857 | 810.7792 | 0.80 |
| **T** | 5 | b5 | 2- | 777.594 | 777.5908 | 0.41 | **T** | 17 | y17 | 7- | 765.0577 | 765.0616 | -0.51 |
| **T** | 6 | b6 | 3- | 650.7132 | 650.7102 | 0.46 | **T** | 16 | y16 | 6- | 839.395 | 839.4026 | -0.91 |
| **G** | 7 | a7-G | 2- | 1025.5919 | 1025.5873 | 0.45 | **G** | 15 | y15 | 6- | 786.0679 | 786.0654 | 0.32 |
| **C** | 8 | a8-C | 3- | 798.4024 | 798.399 | 0.43 | **C** | 14 | w14 | 7- | 638.0448 | 638.0427 | 0.33 |
| **T** | 9 | b9 | 4- | 710.8207 | 710.8167 | 0.56 | **T** | 13 | y13 | 5- | 813.4731 | 813.4693 | 0.47 |
| **C** | 10 | a10-C | 4- | 754.8115 | 754.8091 | 0.32 | **C** | 12 | x12 | 5- | 765.0577 | 765.0512 | 0.85 |
| **T** | 11 | a11 | 4- | 862.5809 | 862.5757 | 0.60 | **T** | 11 | w11 | 5- | 707.6523 | 707.6486 | 0.52 |
| **T** | 12 | b12 | 5- | 757.4689 | 757.4658 | 0.41 | **T** | 10 | y10 | 4- | 780.8244 | 780.8209 | 0.45 |
| **C** | 13 | a13-C | 5- | 792.6635 | 792.6598 | 0.47 | **C** | 9 | y9 | 4- | 700.8193 | 700.8151 | 0.60 |
| **T** | 14 | | | | | | **T** | 8 | y8 | 3- | 833.0214 | 833.0814 | -7.20 |
| **T** | 15 | b15 | 6- | 788.5617 | 788.5653 | -0.46 | **T** | 7 | y7 | 4- | 568.5426 | 568.5393 | 0.58 |
| **C** | 16 | a16-C | 6- | 817.8979 | 817.8936 | 0.53 | **C** | 6 | w6 | 4- | 488.5368 | 488.5335 | 0.68 |
| **T** | 17 | | | | | | **T** | 5 | w5 | 3- | 550.0422 | 550.0393 | 0.53 |
| **T** | 18 | | | | | | **T** | 4 | w4 | 3- | 443.3664 | 443.3649 | 0.34 |
| **G** | 19 | c19 | 8- | 762.1806 | 762.1762 | 0.58 | **G** | 3 | y3 | 1- | 916.1448 | 916.1419 | 0.32 |
| **C** | 20 | a20-C | 8- | 774.4352 | 774.4308 | 0.57 | **C** | 2 | w2 | 1- | 667.0582 | 667.0564 | 0.27 |
| **G** | 21 | | | | | | **G** | 1 | w1 | 1- | 362.0335 | 362.0329 | 0.17 |

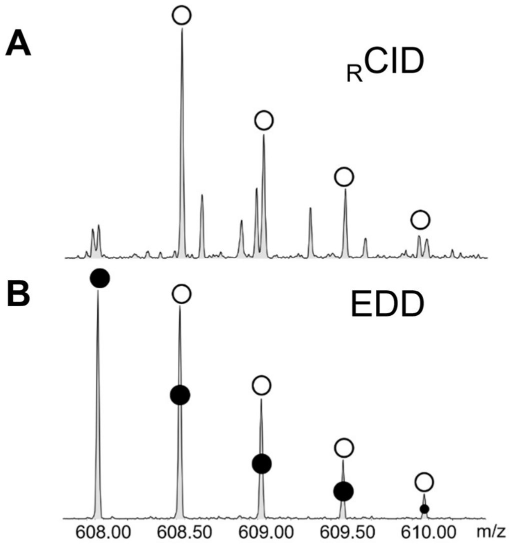


**Figure S4: Isotopic distribution of the $a_4^{2-}$ fragment ion** A) $_R$CID showing closed-shell isotopic distribution and B) EDD showing a mixture of open and closed-shell isotopic distribution. Black circles indicate radical ions, while open circles indicate closed-shell ions. The two species differ by one hydrogen mass divided by the charge state (1.008 Da/z).

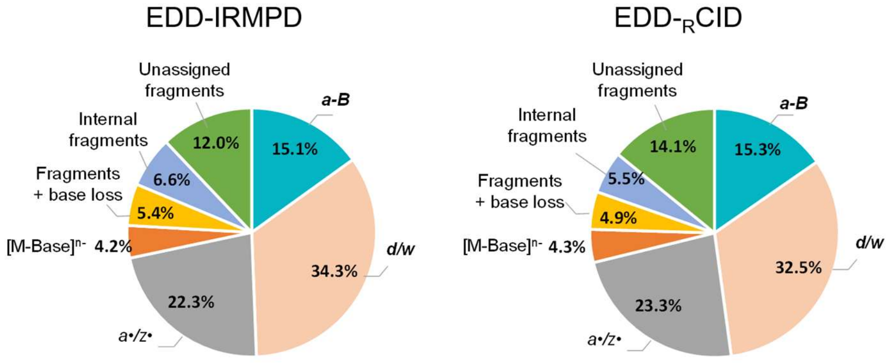


**Figures S5:** Comparison of the fraction of peaks assigned to each fragment category for EDD performed on $[M\text{-}9H]^{9-}$ DNA, followed by IRMPD or $_{R}$CID on re-isolated $[M\text{-}8H]^{8-\bullet}$. All fragment assignments are provided in supporting information Table S8 (separate Excel file). Activation energies: EDD was performed using 30 eV electron kinetic energy and 50 ms irradiation time. Subsequent vibrational activation was achieved using 4.5 ms IRMPD (9.3 μm, 50% power) or 270 mV for $_{R}$CID.

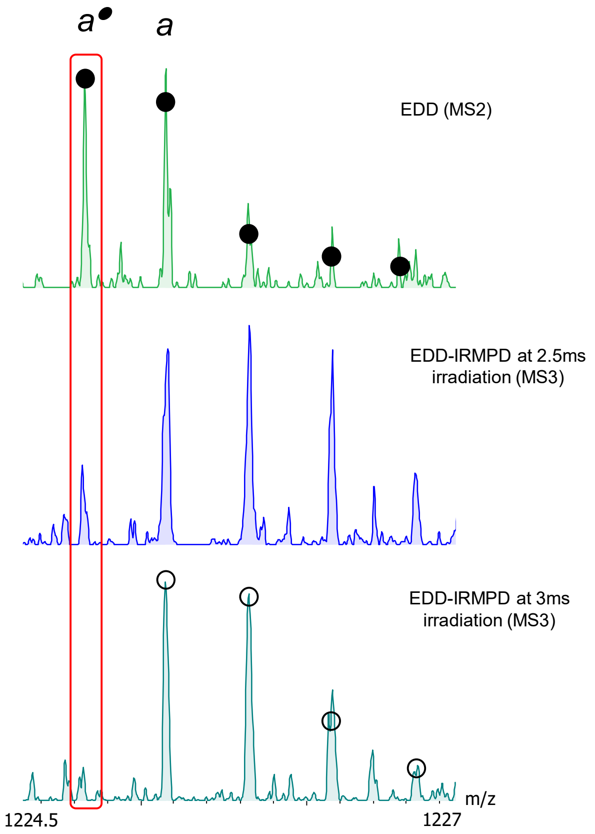


**Figures S6:** Effect of IRMPD irradiation time on the abundance of *a*• radical fragment generated by EDD of the 46mer RNA $[M\text{-}8H]^{9-}$. Black and white circles correspond to theoretical isotopic distributions for radical and closed-shell *a* fragment respectively. With EDD alone ($MS^2$), we almost have full radical $a^{\bullet}$ ions. Upon activation with IRMPD at 2.5 irradiation, we have a mixture of radical and closed-shell *a* fragments. At higher energies (3 ms IRMPD), the isotopic distribution corresponds to a full-closed-shell ion.

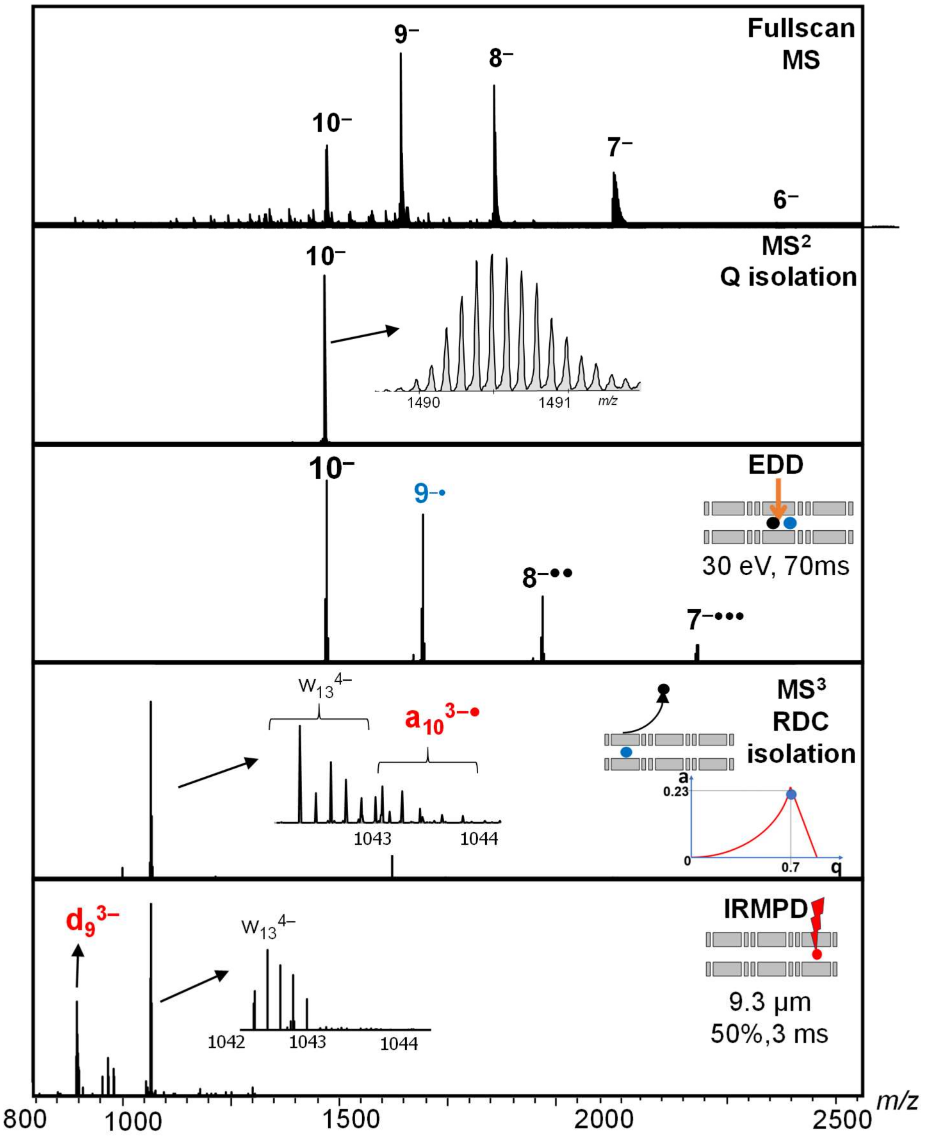


**Figures S7:** MS$^3$ dissociation of radical fragment into a closed-shell ion. From the fullscan MS spectrum of the 46-mer RNA UCUAAGUAAAAUUGGGUGGGUGGGUGGGUUUGUCGGUAUCUCGGUU, the charge state 10- (m/z 1490.9) is isolated using the quadrupole. The ions are accumulated in section Q5 for 250 ms and submitted to EDD. 70 ms of irradiation with electron of kinetic energy 30 eV is used to produce radicals ($9^{-\bullet}$, $8^{-\bullet\bullet}$, $7^{-\bullet\bullet\bullet}$). The radical $a_{10}^{3-\bullet}$ (m/z 1043.15) is isolated from the EDD spectrum in a MS$^3$ stage using resolving DC by changing the drive frequency of the Omnitrap. It was actually co-isolated with a *w* ion, but upon irradiation for 3 ms (IRMPD, 9.3 µM, 50% power), $a_{10}^{3-\bullet}$ disappears and the first resulting fragment is a $d_9^{3-}$.

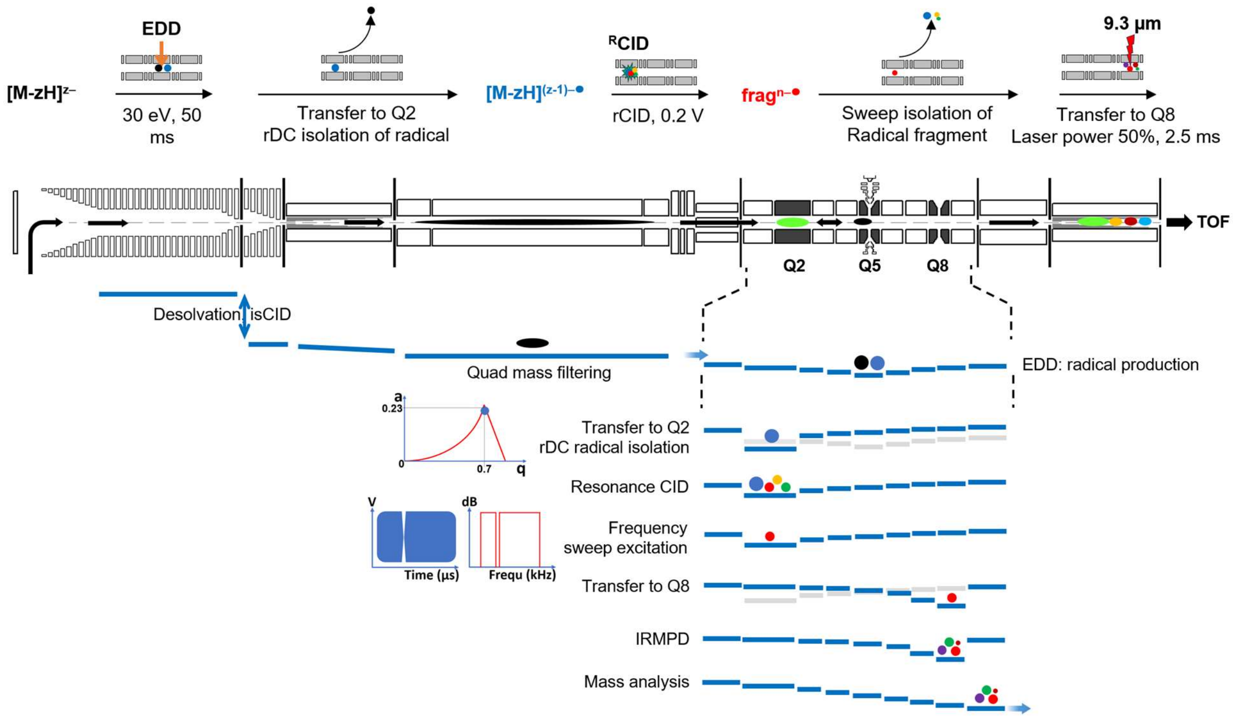


**Figures S8:** Schematic representation of the **MS$^4$** experiment reported in main text Figure 7 and supplementary Figure S10. **MS$^2$**: the precursor ion is selected using the quadrupole and accumulated in Q5. The ion undergoes EDD for 50 ms and the resulting ions are transferred to Q2. **MS$^3$**: in Q2, the radical ion resulting from a single electron loss is isolated using resolving DC and submitted to resonance CID, producing both closed shell and radical fragments. **MS$^4$**: One of the radical fragment is re-isolated using a frequency sweep excitation with a notch corresponding to its secular frequency and then submitted to resonance CID. The fragment ions are transferred toward the mass analyzer. The complete sequence takes 350 ms when using an accumulation time of 80 ms before the EDD process. The blue lines schematically represent the DC voltages applied to the different sections of the Omnitrap. The light gray lines represent the voltages from the previous step for ease of visualization.

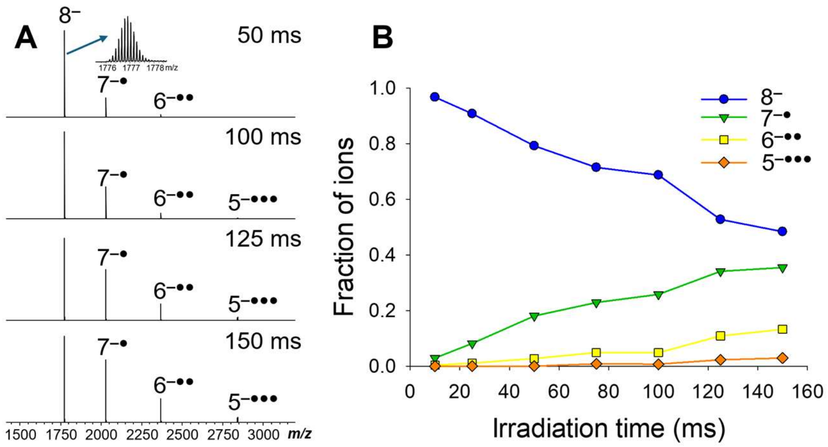


**Figure S9:** Influence of the EDD irradiation time on the 46-mer DNA $[M-8H]^{8-}$. The kinetic energy of the electron is 30 eV.

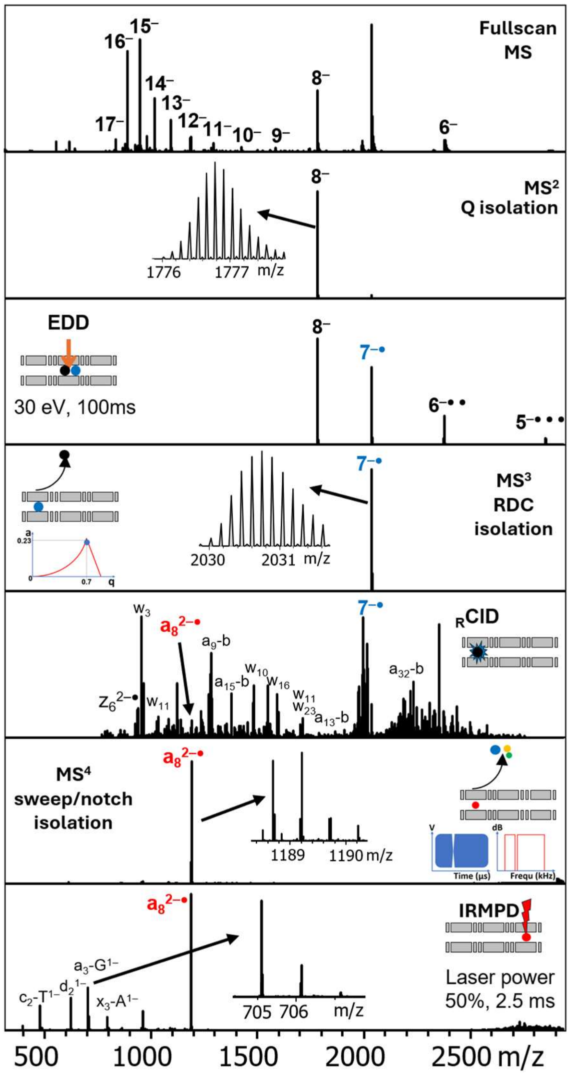


**Figure S10: MS⁴ to elucidate the fate of an *a*• radical fragment from the 46-mer DNA.** From the full scan MS spectrum, the charge state 8– (m/z 1776.9) is isolated using the quadrupole. A description Q1—Q9 DC gradient programming for each step is provided in Supporting Information Figure S8. The ions are accumulated in Q5 for 100 ms and submitted to EDD. 150 ms of irradiation with electron of kinetic energy 30 eV is used to produce radicals (the dependence of radical product ion yields on the irradiation time is shown in Supporting Information Figure S9). The radical $7^{-\bullet}$ (m/z 2030.7) is isolated in a $MS^3$ stage using resolving DC by changing the drive frequency of the Omnitrap. The ions then undergo resonance CID during 10 ms (0.2 V). For the $MS^4$ stage, the radical fragment $a_8^{2-\bullet}$, which is a very minor product in $MS^3$, is re-isolated using a sweep of frequencies including a notch corresponding to its secular frequency. After transfer to section Q8, the radical fragment is irradiated for 2.5 ms (IRMPD, 9.3 µm, 50% power). The resulting fragments are a variety of closed shell ions. The total duration of the sequence is 350 ms, and the reported $MS^4$ spectrum was recorded in 3 minutes.
